# A tyrosine kinase protein interaction map reveals targetable EGFR network oncogenesis in lung cancer

**DOI:** 10.1101/2020.07.02.185173

**Authors:** Swati Kaushik, Franziska Haderk, Xin Zhao, Hsien-Ming Hu, Khyati N. Shah, Gwendolyn M. Jang, Victor Olivas, Shigeki Nanjo, John Jascur, Vincent B. Masto, Daniel Ciznadija, Ido Sloma, Emilie Gross, Scott L. Weinrich, Jeffery R. Johnson, Trever G. Bivona, Nevan J. Krogan, Sourav Bandyopadhyay

## Abstract

Signaling networks balance the activities of many physically interacting proteins and perturbations to this network influence downstream signaling, potentially leading to oncogenic states. Using affinity purification-mass spectrometry we defined this network for all 90 human tyrosine kinases revealing 1,463 mostly novel interactions between these key cancer proteins and diverse molecular complexes. Modulation of interactor levels altered growth phenotypes associated with corresponding tyrosine kinase partners suggesting that tumors may alter the stoichiometries of interactors to maximize oncogenic signaling. We show that the levels of EGFR interactors delineates this form of network oncogenesis in 19% of EGFR wild-type lung cancer patients which were mostly otherwise oncogene negative, predicting sensitivity to EGFR inhibitors in vitro and in vivo. EGFR network oncogenesis occurs through mechanistically distinct network alleles often in cooperation with weak oncogenes in the MAPK pathway. Network oncogenesis may be a common and targetable convergent mechanism of oncogenic pathway activation in cancer.

**HIGHLIGHTS:**

- A human tyrosine kinome protein interaction map reveals novel physical and functional associations.
- Dependence on oncogenic tyrosine kinases is modulated through perturbation of their interactors.
- EGFR network oncogenesis in up to 19% of EGFR wild-type lung cancers is targetable.
- EGFR network oncogenesis cooperates with weak oncogenes in the MAPK pathway.

## INTRODUCTION

Tumors often exhibit mutations in functionally related genes suggesting a limited number of pathways are sufficient to cause tumorigenesis (Sanchez-Vega et al., 2018). However, analysis of tumor genomes have revealed that many tumors lack mutations in such pathways including most notably oncogene negative lung and triple negative breast cancers (Cancer Genome Atlas Research Network, 2014; Shah et al., 2012; Koboldt et al., 2012). In order to reconcile this apparent paradox it is useful to consider that there are multiple ways to dysregulate oncogenic proteins with mutation being the simplest to measure with current technologies (Yaffe, 2013). Other mutation-independent routes of oncogenesis have been described (Bild et al., 2006; Popovici et al., 2012) albeit through mechanisms that remain unclear but could include expression of ligands (Fujimoto et al., 2005; Singh and Harris, 2005; Wu et al., 2007) or loss of factors involved in protein endocytosis and degradation (Bache et al., 2004). While such regulation is multifactorial and an aggregate over largely unknown inputs, proteins do not function in isolation but are rather constrained by physical association with other proteins in its interaction network, i.e. its network context. Hence, elucidating the network context of an oncogene may aid in understanding the pathogenesis of cancers with otherwise unknown disease drivers.

Tyrosine kinases are among the most frequently altered gene families in cancer. The human genome encodes 90 tyrosine kinases and despite their biological and therapeutic importance their unique functions in normal development and diseases such as cancer remain poorly understood (Robinson et al., 2000; Gschwind et al., 2004). They are the predominant class of cancer drug targets, although only roughly 5% of cancer patients match to current indications for available targeted therapy (Marquart et al., 2018). For some drugs that have been investigated broadly, the molecular basis for anti-tumor responses cannot be explained by the presence of a mutation in the target or pathway as biomarker (Ferté et al., 2010; Hirai et al., 2017; Jazieh et al., 2013; Krejci et al., 2011; Loboda et al., 2010; Mirza et al., 2016; Moroni et al., 2005; Shao et al., 2015; Swisher et al., 2017; Tian et al., 2013). In addition, most patient responses to targeted therapies are not durable leading to eventual drug resistance. Comprehensive knowledge of physical interactors for this important gene family could aid in understanding mechanisms of oncogene regulation and consequently delineate new factors important for mediating drug sensitivity and resistance.

One approach to delineate pathway components is through the systematic assembly of protein-protein interaction (PPI) maps (Krogan et al., 2006; Sowa et al., 2009; Breitkreutz et al., 2010; Behrends et al., 2010) Many complementary technologies to map PPIs have been developed including yeast two-hybrid and proteomic approaches (Rual et al., 2005; Hein et al., 2015; Huttlin et al., 2017, 2017; Yao et al., 2017a). Coverage of tyrosine kinases using such experimental approaches has been minimal and is largely biased towards a small number of tyrosine kinases that are most intensely studied. Physical interactions are often also co-functional (Bandyopadhyay et al., 2010) and network maps can provide insights into associations with distinct cellular complexes (Stuart et al., 2006; James et al., 2009), unexpected cellular localizations (Ni et al., 2001; Carpenter, 2003; Carpenter and Liao, 2013) and transactivation partners (Paul and Hristova, 2019). Therefore, a systematic map of interactions associated with this important class of proteins may reveal new and previously unexplored biology outside of canonical signaling cascades.

We systematically mapped physical associations with all 90 human tyrosine kinases by experimentally mapping protein-protein interaction networks using affinity-based proteomics. Analyses of this interactome revealed novel associations of tyrosine kinases with distinct protein complexes reflective of broadly diverse roles in cellular signaling. Mapping chemo-genetic interactions revealed that many of the interactors were also functional and that their perturbation could modulate dependence on oncogenic tyrosine kinase partners. This finding led us to test the hypothesis that measurement of the stoichiometries in protein interaction partners could reveal oncogene dependence and activation state in cancer cells. Applied to the EGFR network, we identified a network-mediated form of oncogenesis that occurs in many otherwise oncogene negative cancers and also co-occurs specifically with weak oncogenic alleles of regulators of the MAPK pathway such as NF1 and BRAF class 3 mutations. This work provides a complete and unbiased interaction map for tyrosine kinases and a new approach to leverage such data to elucidate mechanisms of oncogene addiction.

## RESULTS

### A physical interaction map of human tyrosine kinases

To identify physically associated proteins we performed affinity purification and mass spectrometry on all 90 human tyrosine kinases. Kinases were expressed with a c-terminal 3xFLAG tag by transient transfection into HEK293T cells (Figure 1A). After confirmation of expression, each tyrosine kinase was immunoprecipitated using an anti-FLAG antibody and co-immunoprecipitated proteins were eluted, trypsinized and subjected to mass spectrometry. We performed 419 such affinity purification – mass spectrometry (AP-MS) experiments using at least three biological replicates for every tyrosine kinase (Figure S1A). Overall, we observed an average correlation of 0.7 among replicate experiments (Figure S1B) and 50% average reproducibility in proteins identified (Figure S1C).

**Figure 1:**
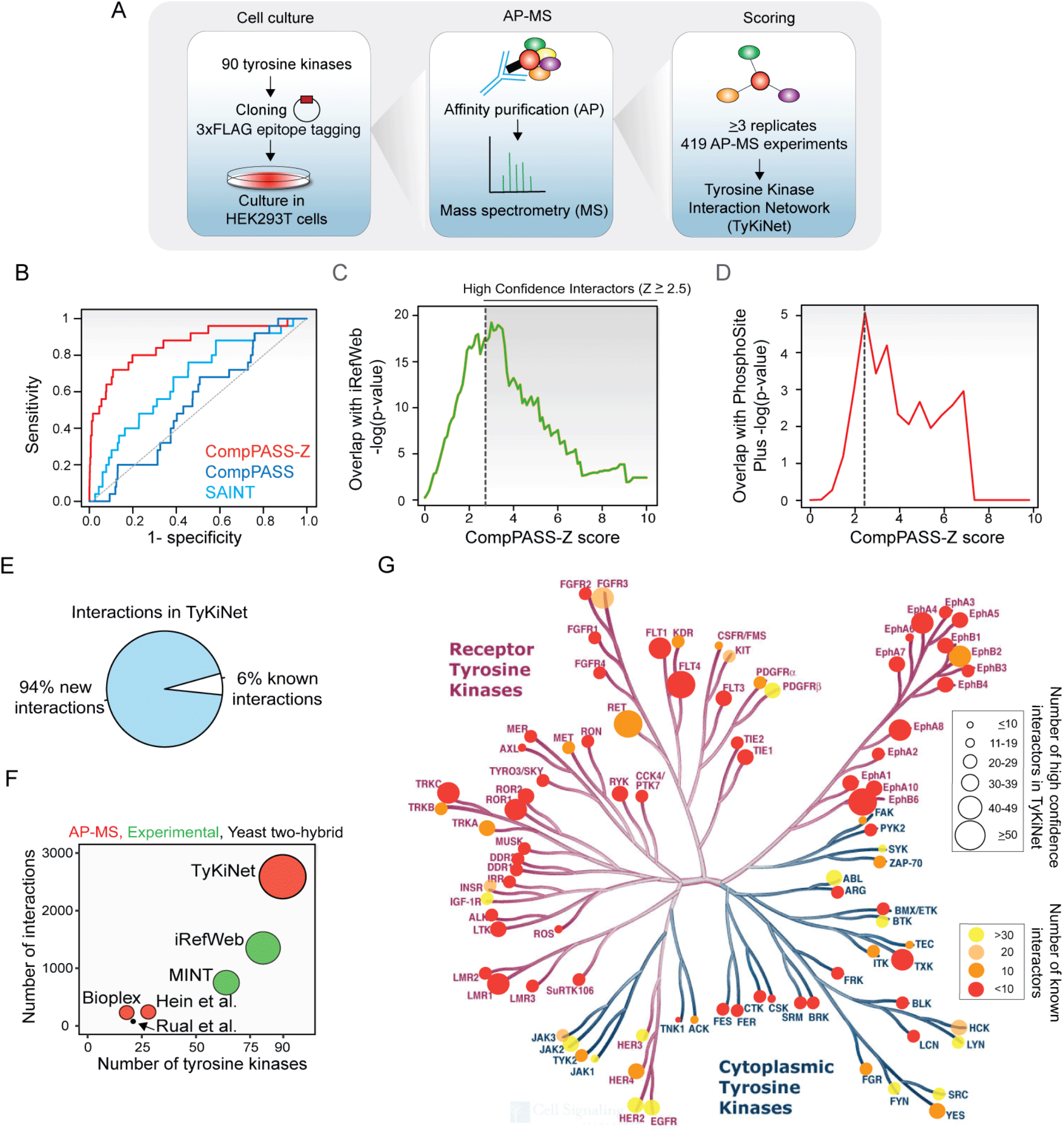
Generation of unbiased tyrosine kinase interactome map. **(A)** Pipeline of Affinity-Purification and Mass Spectrometry (AP-MS) experiments to generate tyrosine kinase interactome in HEK293T cells. **(B)** ROC analysis of enrichment of known tyrosine kinase interactions from HumanNet with AP-MS data scored using CompPASS, CompPASS-Z and SAINT scoring methods. Gray line depicts random prediction. **(C)** Enrichment of known interactions from the iRefWeb database at different CompPASS-Z cutoffs. P-values of enrichment were calculated using hypergeometric test. Black dashed line indicates cutoff used to include high (≥2.5) confidence interactors. **(D)** Enrichment of known substrates of tyrosine kinases from the PhosphoSitePlus database at various CompPASS-Z scores. Dashed line indicates a score of 2.5. P-value of enrichment was calculated using hypergeometric test. **(E)** Fraction of novel and known interactions (iRefWeb, Bioplex, MINT database) of tyrosine kinases found in this study. **(F)** Comparison of number of tyrosine kinase interactors found in different datasets. Red, green and black bubbles indicate AP-MS studies, experimentally identified and yeast two-hybrid studies respectively. **(G)** Number of known and new interactions overlaid onto a tyrosine kinase sequence similarity tree. Size of the bubble represents the number of interactors identified for every tyrosine kinases in TyKiNet. Color represents the number of associated interactions found in the iRefWeb database.

We performed computational analysis to distinguish bonafide from non-specific interactions in the AP-MS data. We evaluated two published methods for their ability to recapitulate known interactions from the raw mass spectrometry data, Comparative Proteomics Analysis Software Suite (CompPASS) (Sowa et al., 2009) and Significance Analysis of Interactomes (SAINT) (Breitkreutz et al., 2010). As the number of AP-MS experiments carried out for every bait varied (Figure S1A), we modified the CompPASS algorithm by including a bait specific normalization (CompPASS-Z, see Methods) then compared the performance of these methods in recapitulating known interactions using two different reference protein-protein interaction (PPI) datasets, HumanNet (Lee et al., 2011) and iRefWeb (Turner et al., 2010). Analysis over different AP-MS score thresholds using Receiver Operating Curve (ROC) analysis indicated that ComPASS-Z distinguishes known interactions with better sensitivity and specificity than other methods (Figure 1B, Figure S1D). We calculated the enrichment of known interactions identified at different ComPASS-Z score cutoffs and found a significant overlap with iRefWeb peaking in significance between scores of 2 and 3 (Figure 1C). Similarly, we found significant enrichment of known kinase-substrate interactions, defined in the PhosphoSitePlus database (Hornbeck et al., 2012), at cutoff of 2.5 (Figure 1D). Therefore, we defined the tyrosine kinase network (TyKiNet) as 1,463 interactions with a score ≥2.5 as high confidence interactions and an additional 1,135 interactions with a score between 2 and 2.5 as lower confidence interactors (Supplementary Table 1).

Analysis of resulting high-confidence interactions revealed that the majority (94%) of interactions were novel and not previously reported in the literature (Figure 1E). This network was larger and had broader coverage of tyrosine kinases than other AP-MS, yeast two-hybrid or experimentally determined interaction databases (Figure 1F). TyKiNet contains a median of 15 interactors per kinase and is unbiased in comparison to our existing knowledge of interactors that is heavily focused on more intensely studied kinases such as EGFR, SRC and HER2 (Figure 1G).

### Tyrosine kinases interact with functionally diverse protein complexes

We sought to place kinase interactors into biological context by analyzing them with respect to known protein complexes and pathways. This analysis identified both known and novel associations between 69 tyrosine kinases and a diverse set of pathways and protein complexes (Figure 2). For example, we detected known interactions between ABL1 and members of the WAVE2 complex (Stuart et al., 2006), and between BTK and the PAF chromatin remodeling complex (Figure 2, Figure 3A,B) (James et al., 2009). A number of novel interactions were also identified such as specific interactions between BMX and the COP9 signalosome, ephrin receptors (EPHA2, EPHA4) and the MCM DNA replication complex, PDGFRB and the RNA exosome complex, and SRC and the Ragulator complex and lysosomal v-ATPase (Figure 3A,B). 40 tyrosine kinases interacted with multiple components of individual protein complexes encoded in the CORUM database in a statistically significant manner (Figure 3A, Supplementary Table 2).

**Figure 2:**
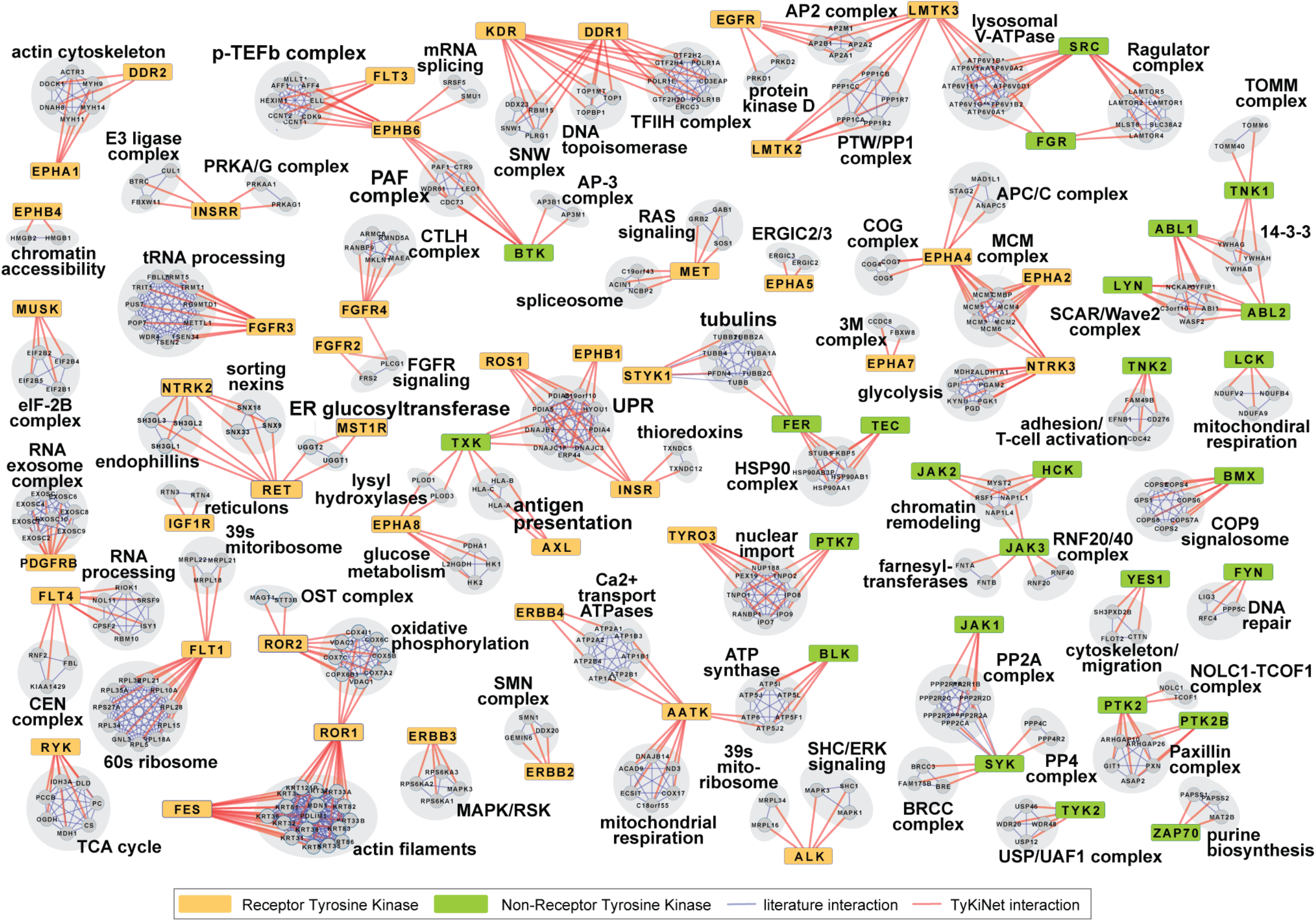
Tyrosine kinases are associated with diverse protein complexes. Network representation of physical interactions in TyKiNet between tyrosine kinases and known protein complexes and functional groups mapped using CORUM and iRefWeb databases. Network connections were identifyed by searching for triangular relationships among tyrosine kinases and multiple members of the complex or pathway (see Methods). Orange and green rectangles indicate receptor and non-receptor tyrosine kinases, respectively. Red interaction edges represent interactions observed in TyKiNet while blue edges indicates known interactions mapped from CORUM or iRefWeb database. Members of the same complex or pathway are grouped in grey.

**Figure 3:**
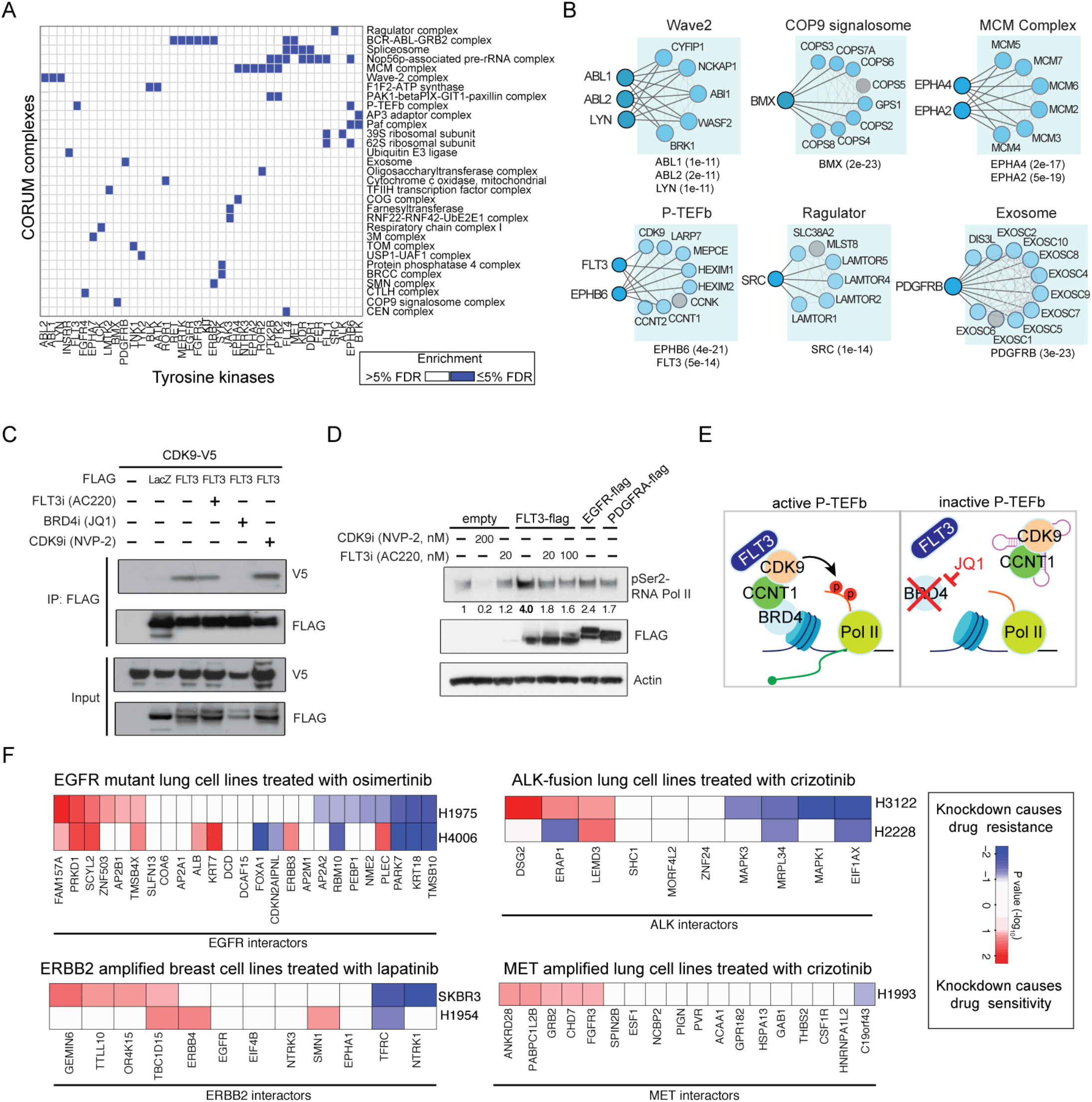
Tyrosine kinase interactome reveals functional relationships in cancer cells. **(A)** Heatmap representation of enrichment of tyrosine kinase interactors with known protein complexes from the CORUM database. Statistical significance of the overlap of interactors of tyrosine kinase with a known protein complex was calculated using hypergeometric test at a 5% FDR cutoff. **(B)** Network representation of protein complex coverage achieved in TyKiNet for selected CORUM complexes. Dark and light blue shades represent bait and prey proteins respectively, while grey circles indicates proteins that were not captured in this study. Black edges from TyKiNet and gray edges from CORUM. P-values of complex coverage based on each tyrosine kinase were calculated as in **(A)**. **(C)** Co-immunoprecipitation of FLT3 tyrosine kinase with CDK9. Cells were exposed to vehicle, 10nM ACC220, 300nM JQ1 or 300nM NVP-2 for 24 hours before harvest. **(D)** Immunoblot of phosphorylation of Ser2 of CTD of the RNA polymerase II after transfection with the indicated empty or flag tagged constructs and untreated or treated with various inhibitors at the indicated dose for 24hours before harvest. Quantification of relative p-Ser2 level relative to untreated empty vector is shown. **(E**) Model of role of FLT3 binding and promotion of the activity of the P-TEFb complex. **(F)** Heatmap representing chemical-genetic interactions involving EGFR, ALK, ERBB2 and MET in cancer cell lines. Interactors of the selected tyrosine kinase oncogene were knocked down using siRNAs in the indicated cell lines. Proliferation after knockdown in the presence and absence of the indicated inhibitor was normalized and compared to determine a P-value of significance (see Methods). Knockdowns were scored for their ability to impart drug resistance (blue) or drug sensitivity (red).

While receptor tyrosine kinases (RTKs) are primarily studied in the context of their localization to the plasma membrane, we observed that many of the protein complexes they interact with are localized in other organelles. Since physical interactions occur when proteins share spatial localization (Schwikowski et al., 2000; Shin et al., 2009), we used the known subcellular localization of interactors encoded in the Gene Ontology to predict the cellular localization of RTKs based on a guilt-by-association principle (Figure S2A). This analysis predicted diverse cellular localizations for RTKs including associations with the nucleus, mitochondria and actin cytoskeleton which we compared to endogenous protein localization determined by immunofluorescence using primary antibodies from the Human Protein Atlas (Thul et al., 2017). We found a localization consistent with our prediction in 19 out of 33 (57%) of cases (Figure S2A). For example, we predicted cytoskeletal localization for DDR2 based on interactions with members of the actin cytoskeleton (DOCK1, MYH11, DNAH8) and which was confirmed by endogenous DDR2 staining in BJ cells (Figure S2B). The majority of RTKs (32/58, 55%) have a predicted nuclear localization based on associations with complexes known to reside in the nucleus (Figure S2A). For example, DDR1 interacts with members of the transcription factor 2H complex (TFIIH) and EPHB6 interacts with the P-TEFb transcription elongation and PAF chromatin remodeling complexes (Figure 2B). Endogenous DDR1 and EPHB6 were both nuclear in cancer cell lines, confirming our prediction (Figure S2B). These data are consistent with previous reports of specific RTKs signaling in the nucleus (Ni et al., 2001; Carpenter, 2003; Carpenter and Liao, 2013), and indicates that many members of this family may have functions in the nucleus. Our systematic analysis indicates that in pathogenic contexts such as cancer where RTKs are often over-expressed (Du and Lovly, 2018) they can take on alternative cellular localizations and associate with functionally distinct protein complexes.

We identified interactions between FLT3 and members of the P-TEFb transcription elongation complex in TyKiNet (Figure 3C) including an interaction between FLT3 and CDK9 (CompPAS-Z score = 2.83), the catalytic subunit of the P-TEFb that phosphorylates Serine 2 of carboxy-terminal domain (CTD) of RNA polymerase II (Marshall et al., 1996). We confirmed the interaction between FLT3 and CDK9 by co-immunoprecipitation (Figure 3C). We next tested if this interaction was dependent on the activation state of FLT3, CDK9 or the entire P-TEFb complex using kinase inhibitors of FLT3 (AC220) or CDK9 (NVP-2) as well as the bromodomain inhibitor JQ-1 which inactivates the P-TEFb by preventing it from binding chromatin via BRD4 (Bartholomeeusen et al., 2012). The interaction between FLT3 and CDK9 was specifically disrupted by JQ1, indicating that the association between FLT3 and CDK9 occurs only in the context of a chromatin-bound, active P-TEFB complex and is not dependent on FLT3 or CDK9 activity (Figure 3C). Further supporting a functional role for this interaction, FLT3 ectopic expression specifically stimulated the activity of the P-TEFb by increasing the phosphorylation of Ser2 of the CTD, which was not observed with expression of control RTKs that do not interact with this complex, EGFR or PDGFRA (Figure 3D). These results indicate that FLT3 promotes the activity of P-TEFb complex when it is bound to chromatin (Figure 3E). We postulate that the numerous novel associations delineated in this interaction map can guide the discovery of other similar functional relationships between tyrosine kinases and protein complexes.

### The tyrosine kinase interactome encodes functional interactions that regulate oncogenic tyrosine kinase dependency in cancer

We next determined if physical interactors could modulate the function of their partner tyrosine kinases. To capture functional interactions between a tyrosine kinase and its physical interaction partner, we systematically measured chemical-genetic interactions using targeted tyrosine kinase inhibitors coupled with knockdown of interactors to measure synthetic enhancement or suppression of drug sensitivity (Hu et al., 2018). We assembled a panel of tyrosine kinase inhibitors that inhibit EGFR (osimertinib), ALK (crizotinib), MET (crizotinib), or ERBB2 (lapatinib) and seven matched RTK driven cancer cell lines harboring EGFR mutation (H1975, H4006), MET (H1993) or ERBB2 amplification (SKBR3, H1954) or EML4-ALK fusion (H3122, H2228). In total, knockdown of 80% of the tested EGFR interactors could significantly modulate sensitivity to the EGFR inhibitor osimertinib in EGFR-mutant lung cancer cell lines (Figure 3E), indicating that loss of interactors can impair (sensitivity) or substitute (resistance) for EGFR signaling. In the case of ERBB2, 66% of interactors modified the response to lapatinib in HER2 amplified breast cancer cell lines. Similar results were also observed when modulating the levels of interactors of ALK in EML4-ALK expressing lung cancer cell lines (70%) as well as interactors of MET in MET amplified cells (33%, Figure 3E). Hence many of the physical interactors defined in this study are also functionally relevant since levels of interactors are important for mediating oncogene signaling.

### Interactor expression defines an EGFR network oncogene in NSCLC

Since knockdown of its interacting partner often modulates kinase signaling we hypothesized that the abundance of interactors in a sample is reflective of the baseline signaling state of a kinase. In this scenario, physical interactors can be considered as allosteric regulators wherein total signaling output is a function of their stoichiometries. Importantly, not all physical interactors are functionally important in this manner and interactors could either positively or negatively influence signaling activity. Together, the levels and activity of proteins in this functional network constitute a form of network mediated protein regulation which we term network activity. Approximating the level of each interactor using gene expression data from a sample, we developed a computational framework to integrate physical interactors of a kinase with tumor transcriptome data in order to determine network activity. Tumors where signaling is known to be maximized were used to train the network activity model by identifying a subset of interactors of a kinase that are differentially expressed, positively or negatively, in cases where the kinase is known to be activated by mutation or amplification. Next, a signed sum of the levels of this subset of interactors is used to quantify the network activity of this kinase in patients (Figure 4A, see Methods). As our initial test case for this approach we interrogated the network of EGFR, an oncogene that is altered by mutation in approximately 12% of lung adenocarcinomas (LUAD) and when mutated is associated with sensitivity to EGFR tyrosine kinase inhibitors (Pao et al., 2004; Lynch et al., 2004; Cancer Genome Atlas Research Network, 2014). We identified physical interactors of EGFR derived from both TyKiNet and literature sources and integrated them with RNA-seq data from 576 TCGA LUAD patient samples. 26 EGFR physical interactors were differentially expressed (FDR<5%) in samples harboring known activating mutations in EGFR (L858R and exon 19 deletion) (Figure 4B). Expression of interactors in each sample was used to calculate EGFR network activity (NA) and permutation analysis and information maximization criteria were used to define tumors that were EGFR network activity positive (EGFR^n+^, NA≥5) (Figure S3A, Supplementary Table 3). At this cutoff, 95% of tumors with canonical EGFR activating mutations were classified EGFR^n+^ (Figure 4B).

**Figure 4:**
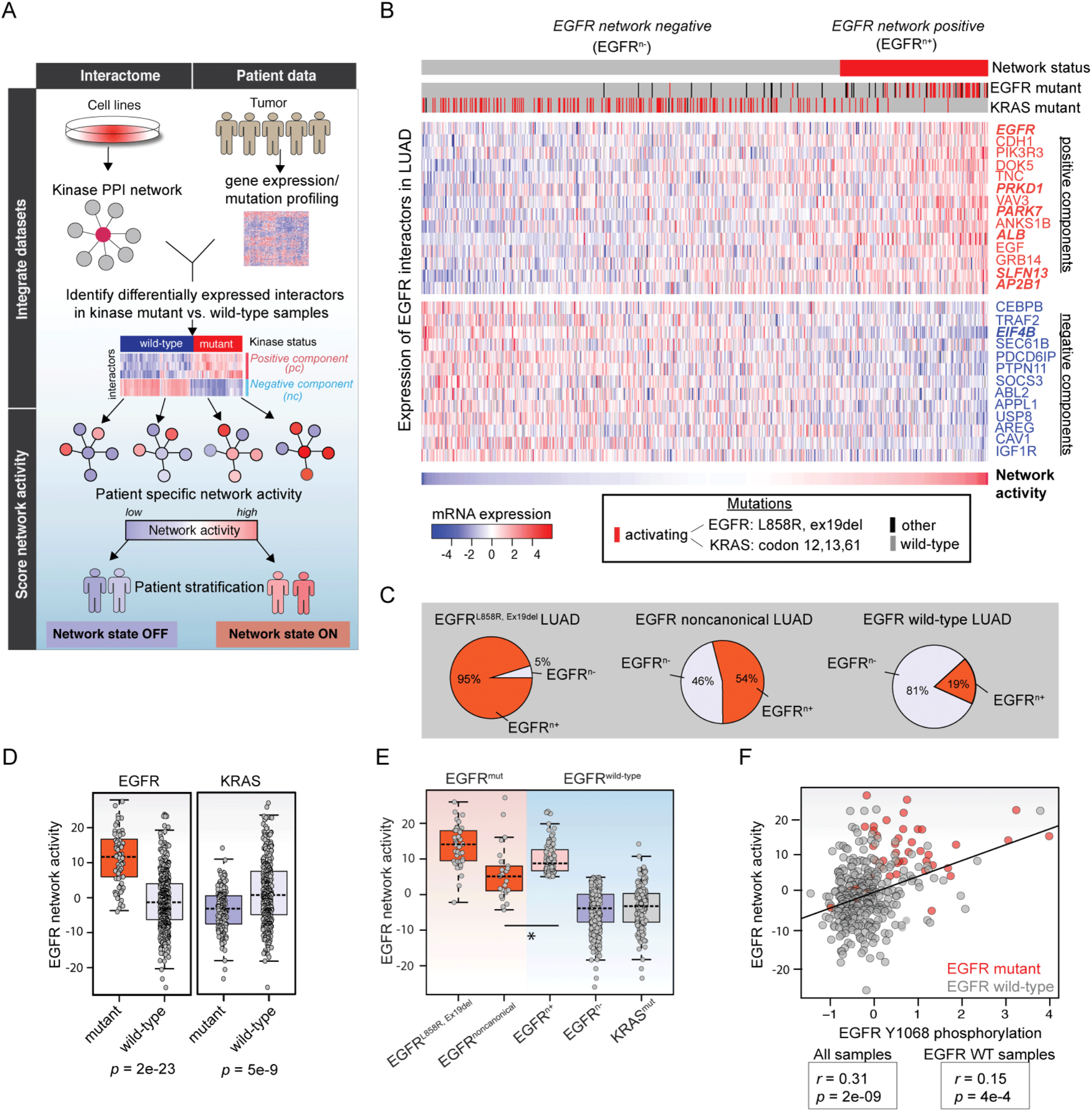
Tyrosine kinase interactome enables identification of EGFR network activity in lung adenocarcinoma. **(A)** Algorithm to map network activity by integration of physical interactors and gene expression data from TCGA. For a given kinase, expression of its interactors was compared between mutant and wild-type samples to identify positively or negatively differentially expressed interactors (5% FDR). In a single sample the network activity is calculated by a signed sum of all such interactors (see methods). **(B)** Heatmap of network activity and constituent EGFR interactors in lung adenocarcinoma patients (LUAD). Columns represent LUAD patients (n=576) and rows represent expression of interactors of EGFR that are either positively or negatively associated with EGFR activity. Samples are sorted from network activity low to high. The mutation status of EGFR and KRAS in each sample is shown above. EGFR interactors from TyKiNet are in bold. **(C)** The fraction of samples with EGFR canonical (L858R or Ex19del) or non-canonical mutation or otherwise EGFR wild-type and EGFR network positive (EGFR^n+^). **(D)** Comparison of EGFR network activity between EGFR and KRAS wild-type versus mutant samples. **(E)** Comparison of EGFR network activity between various groups based on EGFR or KRAS mutation and network activity scores. * = p<0.001 by rank sum test. **(F)** Comparison of EGFR network activity with pEGFR Y1068 levels in LUAD samples. EGFR mutant samples are shown in red. Correlation and p-value are based on Spearman rank correlation. P-values calculated using two-tailed Wilcoxon rank-sum test in (D) and (E). Boxes represent the inter quartile range, and whiskers indicate 1.5 times the interquartile range.

Non-canonical mutations in EGFR are not as recurrent and it is unclear which of these mutations are functional versus non-functional (Berger et al., 2016; Kohsaka et al., 2017; Ng et al., 2018). Over half (54%) of tumors harboring non-canonical EGFR mutations were also EGFR^n+^ which was more likely than background indicating that many non-hotspot mutations in EGFR may also be activating (Fisher Exact test p=0.002, Figure 4C). Non-canonical EGFR mutations in the kinase domain (Wilcoxon rank-sum test p=5.3e-23), receptor L domain (RL) (p=0.007) and growth factor receptor domain IV (GFIV) (p=0.004) domains had significantly higher network activity than EGFR wild-type samples (Figure S3B). Tumors with oncogenic driver events downstream of EGFR such as mutations in KRAS had lower NA scores and generally were classified as network negative (EGFR^n−^) (KRAS mutant vs. wild-type, Wilcoxon rank-sum test p=5e-09, Figure 4D).

We observed that 19% of EGFR mutation wild-type samples were network positive (EGFR^wt,n+^) with network activity scores near that of canonical EGFR-mutant samples and higher than that of those with non-canonical EGFR mutations (Figure 4C,E). When active, EGFR is auto-phosphorylated at Y1068, a docking site for GRB2 that links EGFR and RAS signaling (Batzer et al., 1994; Rojas et al., 1996). EGFR network activity scores in tumors were significantly correlated with phospho-EGFR Y1068 levels in tumors (spearman r=0.31, p=2e-9), which was also true when the analysis was restricted to only EGFR wild-type samples (spearman r=0.15, p=4e-4, Figure 4F). Hence, EGFR^n+^ tumors are prevalent in EGFR wild-type NSCLC where they display evidence of EGFR hyper-activity.

We next determined if EGFR network activity was predictive of response to EGFR inhibition. First, we evaluated 40 NSCLC cell lines and found that network activity scored from baseline gene expression was significantly correlated with phospho-EGFR levels (spearman r=0.48, p=4e-4) and proliferation in response to the EGFR inhibitor erlotinib (spearman r=0.3, p=0.05) and the dual EGFR/HER2 inhibitor lapatinib (spearman r=0.45, p=0.003) (Figure S3C-E). Second, we investigated the in vivo response of 25 NSCLC patient derived xenograft (PDX) models treated with erlotinib in an n-of-1 mouse clinical trial format (Gao et al., 2015). PDX transcriptome data at baseline was used to score EGFR NA in each model. The only EGFR-mutant model in this cohort expresses a non-canonical EGFR S895I mutation, which also had the highest network activity and demonstrated tumor regression with erlotinib treatment (Figure 5A, Supplementary Table 4). In this cohort 10/25 tumors (40%) demonstrated stable disease and 3/25 (12%) displayed a partial response according to RECIST criteria. EGFR network activity was strongly correlated with the percentage of tumor growth inhibition of each PDX model treated with erlotinib (spearman r=−0.52, p=0.007) and models with responses had a significantly higher network activity than those that had stable disease or progressive disease (p=0.003, Wilcoxon rank-sum test, Figure 5B). There was no correlation between EGFR expression and response in this cohort (Figure 5C). Overall the PDX models with the highest network activity (NA>5) had a significantly longer time to progression defined as days taken for tumor doubling in the presence of erlotinib (25.0 versus 13.1 months, p=0.038 log-rank test, Figure 5D). In a prospective analysis, we analyzed a cohort of 17 NSCLC PDX models and scored them for EGFR NA. This cohort had only one EGFR mutant model (CTG-2535, exon19 del) which had the highest EGFR NA as expected (Figure 5E). We selected the PDX with second highest EGFR NA (NA=6.6, CTG-0165), which was classified EGFR^wt,n+^. Erlotinib treatment in this PDX model resulted in significant reduction in tumor volume in comparison to vehicle (p=0.006, two-tailed t-test, Figure 5E).

**Figure 5.**
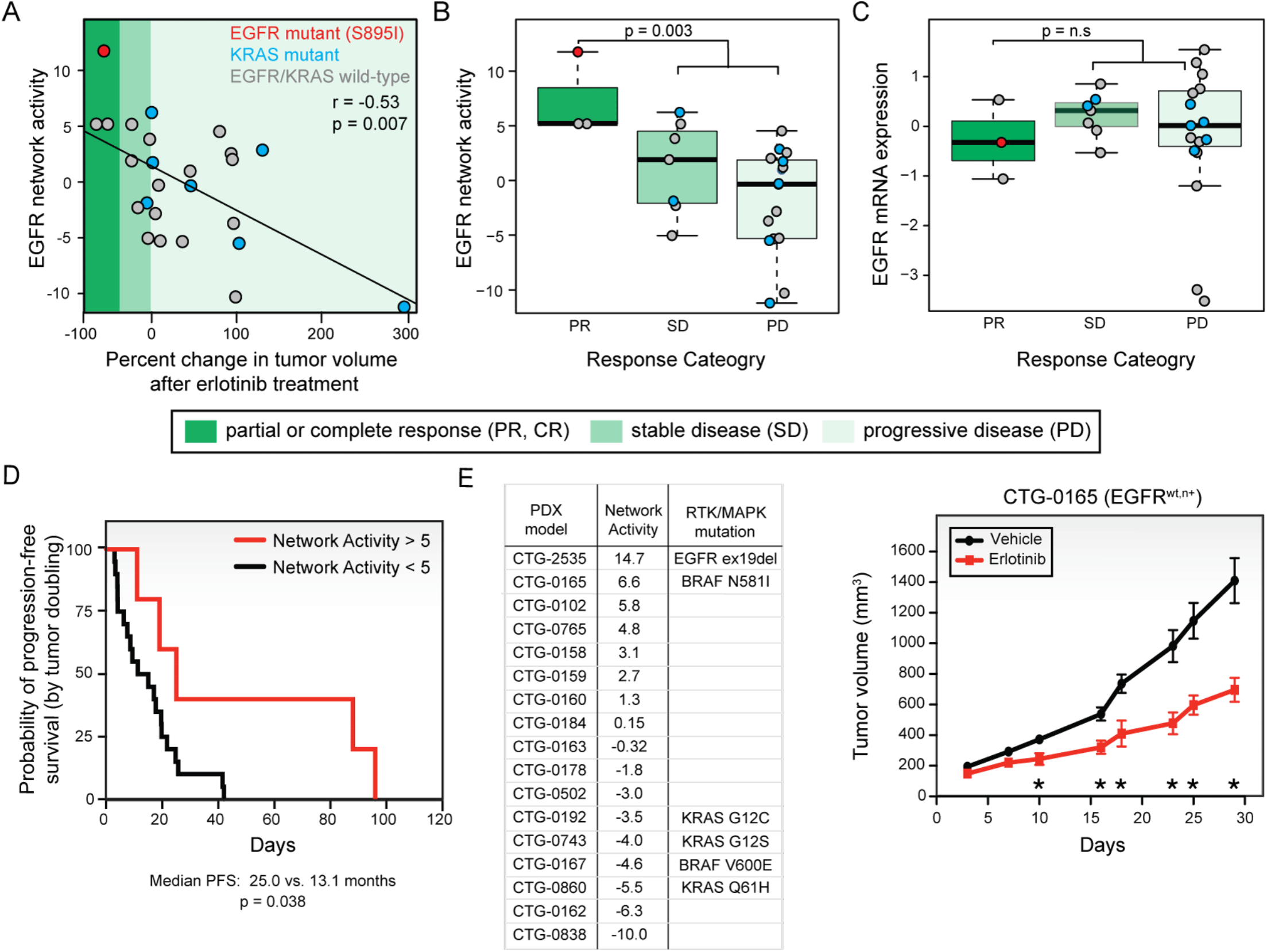
Tyrosine kinase network state predicts targeted therapy response. **(A)** Comparison of baseline EGFR network activity with percent change in tumor volume from the beginning of treatment across 25 different NSCLC PDX models treated with erlotinib in a n-of-1 study from Gao et al. (Gao et al., 2015). P-values are calculated using Spearman correlation coefficient. EGFR and KRAS mutant models are shown in red and blue circles, respectively. Shading corresponds to different response groups. Partial or complete response (PR/CR) corresponds reduction of <−20%, stable disease (SD) corresponds to <30% growth, progressive disease (PD) corresponds to >30% tumor growth compared to initial tumor volume over treatment. EGFR network activity in tumors separated by response group. P-value based on two-tailed Wilcoxon rank-sum test. **(C)** EGFR expression in tumors separated by response group. P-values were calculated using two-tailed Wilcoxon rank-sum test. **(D)** Kaplan-Meier plot of progression free survival of erlotinib treated PDX with EGFR NA>5 (n=5) compared with EGFR NA<5 (n=20). P-value was calculated using log-rank test. **(E)** RNAseq data from an untreated NSCLC PDX cohort was used to score EGFR network activity (n=17). Model with second highest EGFR network activity was EGFR wild-type and network positive (EGFR^wt,n+^) and treated with erlotinib (50 mg/kg/d) and tumor volumes compared to vehicle treatment *p<0.001 two-tailed t-test.

We conclude that in EGFR^wt,n+^ tumors, EGFR network activation and resulting dependency is distinct from mutation or changes in expression of the receptor itself (an effect in *cis*), but rather caused by an activation induced by changes in the balance of proteins interacting with it (*trans* effects). The fact that the EGFR network includes genes that are physically associated with EGFR is critical to the approach, as applying the same methods to generate a signature using a non-EGFR centric network was not as predictive of EGFR phosphorylation in tumor samples, responses to erlotinib in PDX models and EGFR inhibitors in cell lines (Figure S3F). Also compared to expression signatures derived using unbiased machine learning methods the EGFR network approach was more robust in prediction across the same datasets (Figure S3G,H). We conclude that EGFR network activation can occur independent of mutations in EGFR often activating the receptor and driving the growth and proliferation of NSCLCs, a hallmark of oncogenic activity.

### Distinct states reflect alleles of the EGFR network oncogene in NSCLC

The expression patterns of EGFR interactors were largely opposing between EGFR mutant and wild-type samples. Interestingly, the expression level of EGFR interactors among EGFR^wt,n+^ samples was considerably heterogeneous, suggesting the presence of distinct mechanisms of network activation (Figure 6A). To elucidate if there were predominant mechanisms of network activation we applied a data compression technique to condense the profiles of EGFR^n+^ tumors into distinct states using non-negative matrix factorization (NMF) (Figure 6B, S4A) (Kim et al., 2017). EGFR^n+^ tumors could be represented using seven transcriptional components (Figure 6B, Figure S4B-D), which can then be used to hierarchically cluster samples into six distinct network states (Figure 6C, Supplementary Table 5). Each state could be characterized by differential activation or repression of distinct genes in the network. For example, state 4 tumors (12% of EGFR^n+^ samples) often display high levels of EGFR expression as well as the EGFR ligand EGF (Figure S4E). State 5 tumors (13% of EGFR^n+^ samples) harbored highest expression of PIK3R3 an effector bridging EGFR with the downstream PI3K pathway. State 1 tumors had the lowest levels of ABL2, a protein which promotes EGFR internalization by endocytosis (Tanos and Pendergast, 2006). Hence, similar to the concept of distinct oncogenic mutant alleles, we identified six recurrent oncogenic network alleles of EGFR promoting *trans* activation of EGFR through distinct mechanisms.

**Figure 6:**
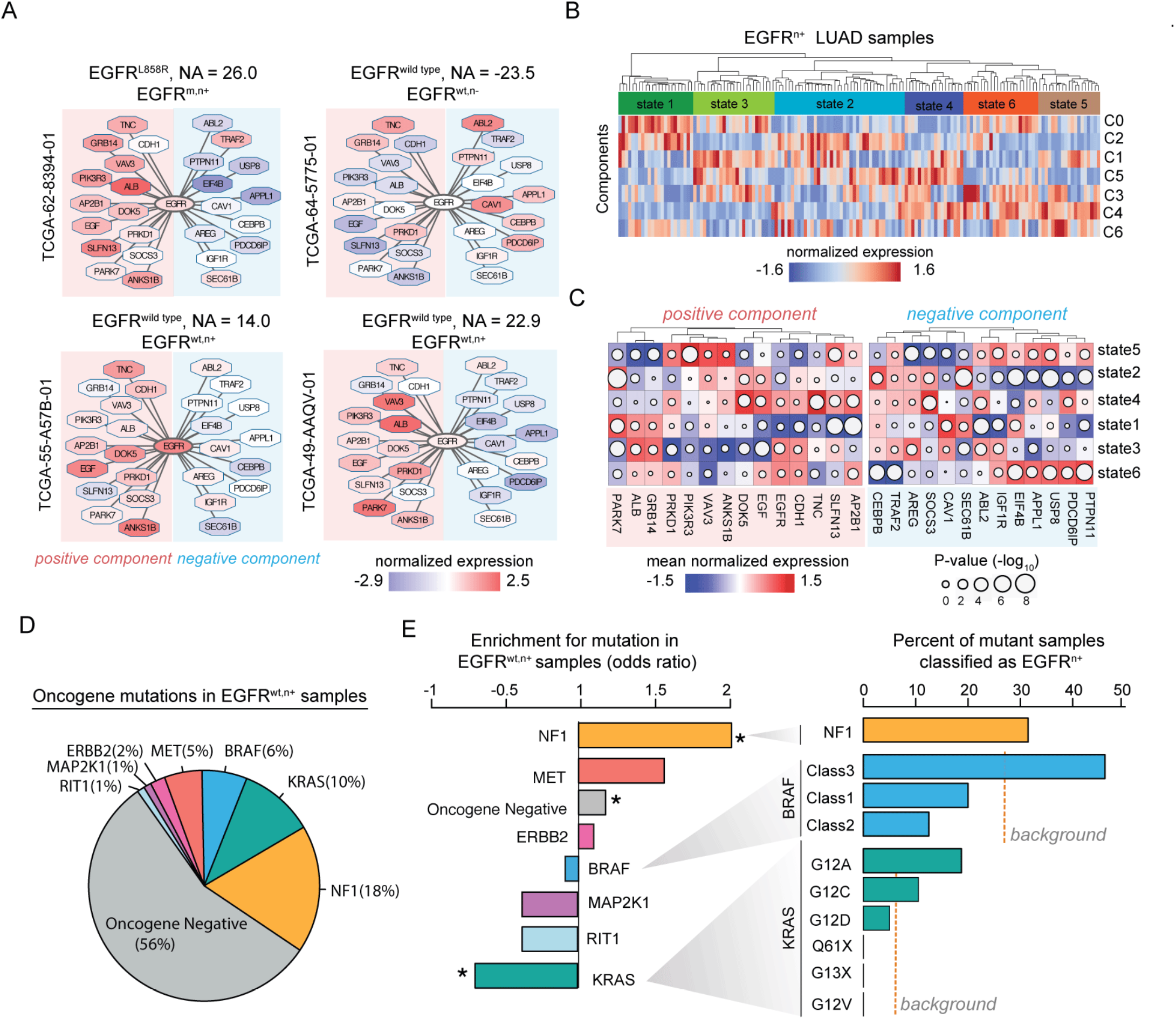
EGFR network state defines distinct mechanism of EGFR activation and is associated with weak oncogenes. **(A)** Representation of EGFR network components in EGFR mutant (L858R), wild-type and EGFR wild-type but network positive (EGFR^wt,n+^) LUAD samples from the TCGA. Node color corresponds to relative mRNA expression of an interactor in a sample. Interactors are separated by positive (left side, red background) and negative (right side, blue background) components. NA = network activity. **(B)** NMF of EGFR network positive samples (EGFR^n+^) indicates seven transcriptional components that cluster into six distinct states. **(C)** Heatmap of the mean expression of interactors of EGFR in six distinct states of EGFR^n+^ patients. Difference in mean expression between samples in each state is shown (color) and significance (size of circle) was calculated using Wilcoxon rank-sum test. **(D)** Occurrence of mutations in known oncogenes and NF1 (Cancer Genome Atlas Research Network, 2014) in EGFR mutation wild-type, network positive (EGFR^wt,n+^) samples (n=94). **(E)** Enrichment of frequency of mutations for specific genes in EGFR^wt,n+^ samples compared to EGFR^n−^ samples. P-value of mutation enrichment was derived from hypergeometric distribution. * = p<0.05. Right barplot represents fraction of all samples with mutation in NF1, or listed alleles of KRAS and BRAF that are also EGFR^n+^. Dashed line indicates the background proportion of all KRAS or BRAF mutants that are EGFR^n+^. X=any amino acid substitution.

### The EGFR network oncogene co-occurs with weak oncogenes in the MAPK pathway

Since co-occurring events in cancer cells may collaborate to promote tumorigenesis, we wondered if mutations that present in EGFR NA high tumors might reflect the tumor selective pressures that result in a requirement for EGFR network oncogenesis. Of all EGFR^wt,n+^ samples, the most common mutation in cancer genes in these tumors was NF1 mutation (18%), followed by KRAS (10%) and BRAF mutations (6%) (Figure 6D). Fifty-six percent were otherwise oncogene negative (Cancer Genome Atlas Research Network, 2014). Although 16% of all LUADs were EGFR^wt,n+^, they constitute 31% of all NF1 mutant cases in this cohort indicating a significant enrichment (p=0.01 by hypergeometric test) (Figure 6E). Inactivating mutations in the RAS-GAP NF1 promote GTP-bound active RAS to activate the MAPK pathway but are generally not sufficient for tumorigenesis (Largaespada et al., 1996; Bollag et al., 1996; Bajenaru et al., 2002). Since different mutant alleles of KRAS and BRAF also variably activate the MAPK pathway (Hunter et al., 2015; Haigis, 2017; Yao et al., 2017b), we tested for selection of distinct oncogene alleles in EGFR^n+^ cases. Codons 12 and 13 encode the major hotspot mutations in KRAS of which G12A and G12C mutations had the highest network activity (p=0.01 one-way anova; Figure S5A). Overall 19% of KRAS G12A and 11% of G12C samples were EGFR^n+^ (Figure 6E). These two alleles of KRAS are among the weakest at this codon due to reduced RAF binding and increased rate of intrinsic hydrolysis (Hunter et al., 2015; Haigis, 2017). BRAF mutant alleles can be categorized into three major classes ranging from strong and monomeric (class 1) to weak and dimeric (class 3) kinase activity (Yao et al., 2017b). 46% of all BRAF class 3 mutant tumors were also EGFR^n+^ compared to 12.5% and 20% for class 2 and class 1 tumors respectively. Hence, mutations that weakly activate MAPK signaling such as in NF1, KRAS G12A/C or BRAF class3 more commonly co-occur with EGFR network oncogenesis in LUAD. While other known cancer associated mutations were not significantly enriched to co-occur across all EGFR^n+^ samples, we observed that mutations in several tumor suppressors were associated with the presence of distinct network alleles of EGFR. For example, although RB1 mutant samples were not enriched in this cohort, 35% of EGFR^n+^ state 5 samples were RB1 mutant (Figure S4F). Similar findings were observed for NOTCH1 (state 1) and RBM10 (state 3) suggesting that EGFR network activation may be fine-tuned based on the nature of the co-occurring secondary mutation.

### Tumors with NF1, KRAS G12A/C, and BRAF class 3 mutations leverage the EGFR network oncogenesis

Although KRAS mutant tumors are currently contraindicated from receiving EGFR targeted therapy our results suggest that some KRAS mutant tumors identified through EGFR network activity may respond to EGFR inhibitors. In the 16 KRAS-mutant NSCLC cell lines available in the Cancer Cell Line Encyclopedia (Barretina et al., 2012), network activity was predictive of response to erlotinib (r=0.62, p=0.01) (Figure 7A). To tested if this trend extended to other EGFR inhibitors we experimentally determined IC_50_ of response to the covalent EGFR inhibitor dacomitinib in a panel of 9 KRAS mutant NSCLC lines and found response was significantly correlated with EGFR network activity (r=−0.79, p=0.012, Figure 7B, Supplementary Table 6). Three cell lines which harbored KRAS mutation (G12A, G12C and G13C) had IC_50s_ below 60nM, a clinically relevant dose given average total plasma exposure of dacomitinib in patients is 161nM (Bello et al., 2013). Since oncogenic mutations in the MAPK pathway cause constitutive signaling, cells harboring such mutations are often thought to be non-responsive to upstream RTK signaling. To test if this was the case we investigated whether EGFR signals to ERK in NF1, KRAS G12A/C and BRAF class 3 mutant tumors. In 6 candidate lung cell lines harboring these genotypes, EGFR suppression with erlotinib led to a reduction of pERK levels similar to that in EGFR-mutant lines and in contrast to that in class1 BRAF (V600E) or ALK-rearranged NSCLC lines whose pERK levels were unresponsive to EGFR inhibition (Figure 7C). Cell lines were sensitive to EGFR inhibition in long term colony formation assays (Figure 7D, Figure S5B). To validate these findings in vivo we treated H358 KRAS G12C xenografted mice with erlotinib and observed significant reduction in tumor volume as compared to vehicle (Figure 7E). These data indicate that EGFR signaling to ERK is functional in tumor cells with high EGFR network activity, including tumor cells harboring mutations in the MAPK pathway, leading to sensitivity to inhibitors of wild-type EGFR.

**Figure 7:**
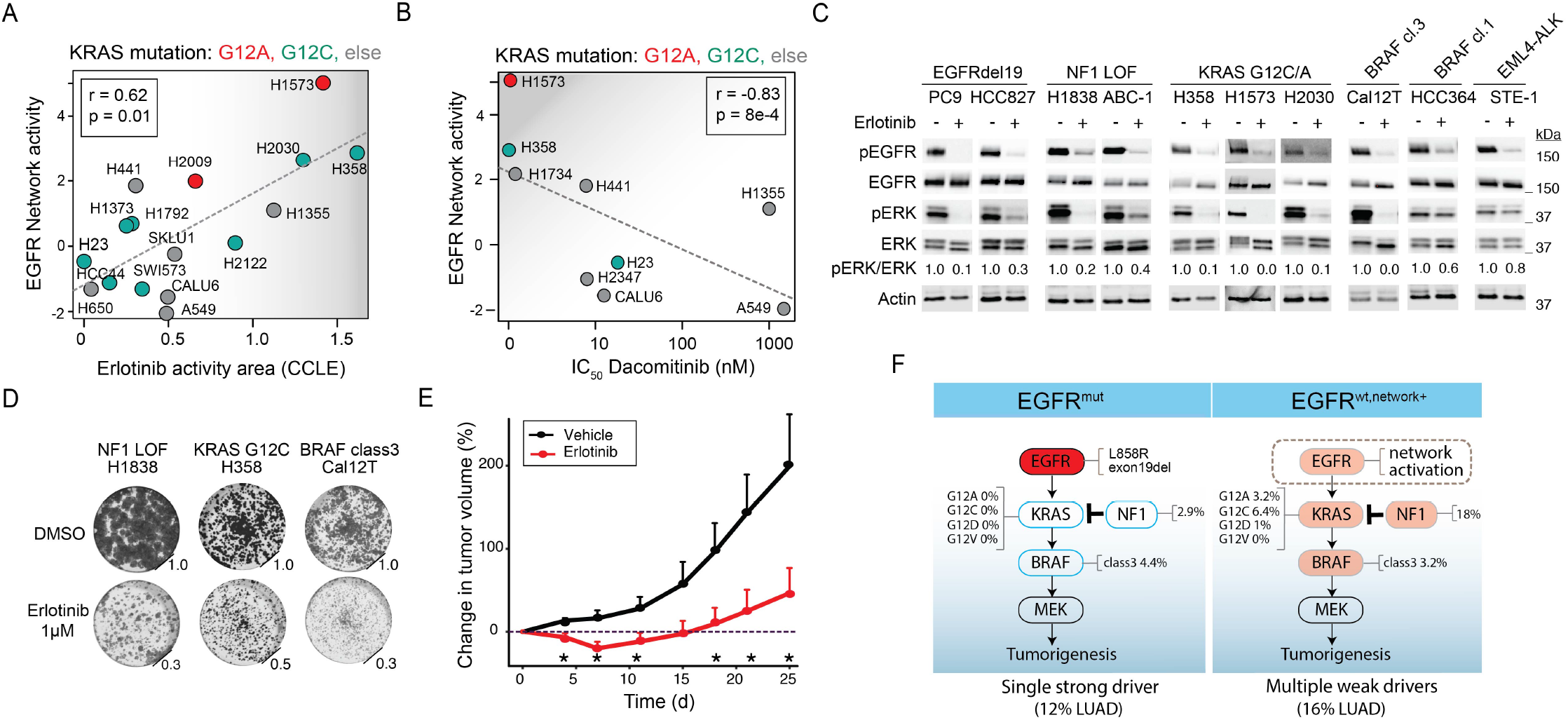
Tumors harboring weak oncogenes are dependent on EGFR signaling. **(A)** Comparison of EGFR network activity (NA) in KRAS mutant NSCLC cell lines with published erlotinib sensitivity (defined as activity area in the CCLE dataset). **(B)** Comparison of dacomitinib sensitivity (IC_50_, determined in this study) with EGFR NA in nine KRAS mutant NSCLC lines. For **(A)** and **(B)** colors represent the type of KRAS mutation and correlation, p-value determined using spearman correlation. **(C)** Immunoblot of pEGFR and ERK levels in lysates from NF1, BRAF-class3 and KRAS G12C and G12A mutant cell lines before and after treatment with erlotinib (1 μm) or 72 hours. PC9 (EGFR del), HCC827 (EGFR del), HCC364 (class1 BRAF) and STE-1 (ALK fusion) cell lines were used as a control. **(D)** Colony formation assay of NF1, BRAF and KRAS mutant lung cancer cell lines grown in the presence or absence of erlotinib for 7 days. **(E)** Growth of H358 (KRAS ^G12C^) mouse xenograft treated with erlotinib (80-100mg/kg/d). * p <0,05, t-statistics. **(F)** A model showing the mutation frequency of NF1, weak oncogenes (class3 BRAF, KRAS G12C, G12A) in patients with EGFR mutant oncogenes and EGFR^wt,n+^ LUAD patients.

Activation of the MAPK pathway by RAS and RAF mutations suppresses EGFR activity and signaling, via negative feedback (Li et al., 2008; van Houdt et al., 2010; Sunaga et al., 2011). Hence it is unlikely that the RAS/MAPK mutations we identified here directly cause EGFR network activation and suggest that EGFR network activation is an independent tumor-promoting event selected for during tumorigenesis. To disprove the null model that mutant KRAS causes EGFR network activation we used H358 NSCLC cells that are KRAS G12C mutant, have high EGFR network activity, respond to erlotinib with an IC_50_ below 1uM and respond to dacomitinib at 9nM indicating that they are EGFR dependent (Figure 7A,B). KRAS G12C inhibitor treatment for 24 hours in H358 cells (Janes et al., 2018; Lou et al., 2019) results in an increase in EGFR network and pathway activity measured by RNAseq (Figure S5C-E) indicating that mutant KRAS does not cause network activation, but rather works to suppress EGFR in these cells. We conclude that the EGFR network oncogenesis is an independent tumor-promoting event (Figure 7F) that is unique from other oncogenic events in the cell that is likely the result of finely tuned selection during tumor development.

## DISCUSSION

The network defined in this study facilitates exploration of the functions of human tyrosine kinases through their protein-protein interactions. In our affinity purification and mass spectrometry approach we used multiple biological replicates and a new metric for scoring which accounts for bait specific variability in contrast to related high throughput approaches (Huttlin et al., 2015, 2017). This resource delineates many new interactions between tyrosine kinases and diverse cellular machinery. Surprisingly, many RTKs interact with proteins that localize most notably in the nucleus suggesting that many RTKs may become aberrantly localized there when overexpressed in pathologic contexts. These data are consistent with reports of nuclear localization of various RTKs (Ni et al., 2001; Carpenter, 2003; Carpenter and Liao, 2013) and indicate that this may be a feature of most members of this family. Furthermore, through interactor focused genetic screens we identified that many of the physical interactors of tyrosine kinases were also functionally important, since the proliferative response of a cell to inhibition of a tyrosine kinase was often modified through the loss of its interaction partner, constituting genetic interaction (Zhang et al., 2016). Practically, interactors may help select drug combinations that may deepen initial tyrosine kinase inhibitor responses or overcome drug resistance.

Tumors often exhibit mutations in distinct but functionally related genes suggesting a limited number of core pathways are sufficient to cause tumorigenesis (Cancer Genome Atlas Research Network, 2014). As a result of this selective pressure we propose that another mechanism to activate a core tumorigenic pathway is through perturbation in the levels of regulators of proteins with oncogenic potential, which we term network oncogenesis. 21% of oncogene-negative lung cancers were EGFR network positive, indicating that network oncogenesis provides a conceptual framework for understanding the pathogenesis of currently untargetable tumors.

We chose EGFR in lung cancer because of the availability of enough patient samples with mutations to train our models and sufficient cell line and PDX models to qualify our approach. As more tumor –omics datasets and models become available, we anticipate this approach will be more broadly applicable. We estimate that the EGFR network oncogenesis occurs in approximately 19% of EGFR wild-type tumors. Historical clinical trials with EGFR TKIs have observed a 6-10% objective response rate in EGFR wild-type tumors (Popovici et al., 2012; Osarogiagbon et al., 2015; Hirai et al., 2017) providing a starting point for patient selection. At present, KRAS mutant tumors are contra-indicated from receiving EGFR TKIs. Surprisingly, many EGFR network positive tumors harbored specific alleles of KRAS, BRAF and mutations in NF1 thus calling for a more nuanced approach to target EGFR in tumors with mutations that activate the MAPK pathway. In support, previous work has shown the importance of pan-ERBB signaling in KRAS-mediated tumorigenesis in mouse models (Kruspig et al., 2018; Moll et al., 2018), exceptional responses to EGFR TKIs have been observed in KRAS G12C mutant lung cancers (Ferté et al., 2010; Krejci et al., 2011) and SHP2 inhibitors which impair RTK signaling to RAS are reported to have activity in many of the same MAPK-related genotypes as in our study (Nichols et al., 2018). We predict that in many cases both network and mutant oncogenes cooperate as the tumor driver. While in these cases single agent EGFR TKIs may inhibit tumor growth, one might anticipate a stronger anti-tumor response in combination with an inhibitor that also targets the mutant oncoprotein directly or downstream such as with KRAS^G12C^ or MEK inhibitors. Our results call for clinical trials testing EGFR TKIs in select NSCLC patients harboring the EGFR network oncogenesis and demonstrate the utility of protein-protein interaction approaches toward realizing network medicine.

## AUTHOR CONTRIBUTIONS

S.K. and S.B. contributed towards study conceptualization. S.K. performed all the data analyses supporting the study. F.H, X.Z, H.-M.H, K.N.S., G.M.J., V.O., S.N., J.J.,V.B.M., D.C., I.S., E.G., S.L.W., designed and performed experiments. J.R.J., T.G.B., N.J.K., S.B. administered the project. S.K. and S.B. composed the original draft and all authors contributed towards manuscript finalization. S.B. supervised the study.

## ACKNOWLEDGEMENTS

We thank Collin Blakley, Eric Collisson, Max V. Ranall, Erik Verschueren, John Von Dollen, UCSF pre clinical core and members of Bandyopadhyay laboratory for helpful discussions and technical assistance. This work was supported by NCI U01CA168370 (S.B.), NIGMS R01GM107671 (S.B.), NCI U54 CA224081 (S.B., T.G.B.) and UCSF Catalyst program (S.B.).

## EXPERIMENTAL PROCEDURES

### Cell culture and reagents

All cell lines were cultured in RPMI supplemented with 10% fetal bovine serum and 1% penicillin/streptomycin. Erlotinib, Lapatinib, AZD9291, TAE684, JQ1, AC220, and Crizotinib was purchased from Selleck Chemicals,. Dacomitinib is available from Sigma. Primary antibodies against phospho-EGFR (#3777, Tyr1068, 1:1000), EGFR (#4267, 1:1000), phospho-ERK (#9101, Thr202, Tyr204, 1:1000), beta-actin (#A2228, 1:2000) and ERK (#9102, 1:1000) were obtained from Cell Signaling Technologies. The antibody against V5 (#SC-81594,1:1000) was purchased from Santa Cruz, Flag (#F1804,1:1000) from Sigma and Phospho-PolI (#5095, Ser2,1:5000) from Abcam.

### Affinity purification

All ORFs were cloned as Gateway entry clones in the pDONR223 vector and subsequently transferred into c-terminal 3xFLAG expression vectors (pcDNA4/TO 3x FLAG). Each vector was confirmed via western blot in HEK293T cells and relative expression levels were determined. All constructs were verified by sequencing. For affinity purification, cells were grown in 15 cm plates and next day transfected with between 3-10 μg of plasmid depending on relative expression levels using calcium phosphate. Each clone was transiently transfected into HEK293T cells. 42 h after transfection, cells were detached and washed with PBS. For Affinity Purification (AP), 2.5×108 cells were induced with 1 μg/ml doxycyclin for 16 h. Cells were lysed in 1 ml cold lysis buffer (50 mM Tris pH 7.5, 150 mM NaCl, 1 mM EDTA, 0.5% Nonidet P40, complete protease inhibitor (Roche) and phosphostop (Roche). Cells were dounced 20x on ice and spun at 2800xg for 20 min. The supernatant was incubated with 60 μl preclearing beads (mouse IgG agarose, SIGMA or Sepharose 4FF, GE Healthcare) for 2 h. The precleared lysate was incubated with 30 μl IP beads over night. FLAG APs were performed with anti-FLAG M2 Affinity Gel (SIGMA) and Strep APs with StrepTactin Sepharose (IBA). The beads were washed 5x with lysis buffer containing 0.05% Nonidet P40 followed by one wash with lysis buffer without detergent. Proteins were eluted with 40 μl 50 mM Tris pH 7.5, 150 mM NaCl, 1 mM EDTA containing either 100μg/ml 3xFLAG peptide (ELIM) and 0.1% RapiGest (Waters), or 2.5 mM Desthiobiotin (IBA). 4 μl of the eluate was analyzed by 4-20% SDS PAGE (Biorad) and silver staining. For co-immunoprecipitations the same procedure was performed on 10^6^ cells and samples were boiled in sample buffer prior to immunoblot.

### Sample preparation for mass spectrometry

For gel-free Mass Spectrometry (MS) analysis 10 μl of the IP eluate were reduced with 2.5 mM DTT at 60°C for 30 minutes followed by alkylation with 2.5 mM iodoacetamide for 40 minutes at room temperature. 100 ng sequencing grade modified trypsin (Promega) was then added to the sample and incubated overnight at 37°C. The resulting peptides were concentrated on ZipTip C18 pipette tips (Millipore) and eluted in a final 20 ul solution of 0.1% formic acid. For gel-based analysis, 20 μl IP eluate was separated by 4-20% SDS-PAGE and stained with GelCode Blue (Thermo Scientific). Each lane was cut into 15 pieces. Each gel piece was diced into small (1 mm^2^) pieces and washed 3x with 25 mM NH4HCO3/50% ACN. Gel pieces were dehydrated and incubated with 10 mM DTT in 25 mM NH4HCO3 and incubated for 1 hour at 56°C. The supernatant was removed and the gel pieces were incubated with 55 mM iodoacetamide and incubated for 40 minutes. Gel pieces were washed with 25 mM NH4HCO3, then 25 mM NH4HCO3/50% can and were then dehydrated. 10 ng/μl trypsin in 25 mM NH4HCO3 was then added to the gel pieces and incubated overnight at 37°C. Finally, peptides were extracted from the gel pieces with 50% ACN/5% formic acid and the solvent evaporated. The final peptide sample was resuspended in 20 μl 0.1% formic acid.

### Mass spectrometry primary data analysis

All samples were analyzed by LC-MS/MS on a Velos Pro dual linear ion trap mass spectrometer (Thermo) equipped with a nanoACQUITY UPLC (Waters) chromatography system and a nanoelectrospray source. 5 μl of each sample was injected onto a a nanoACQUITY Symmetry C18 trap (5 μm particle size, 180 μm x 20 mm) in buffer A (0.1% formic acid in water) at a flow rate of 4 μl/min and then separated over a nanoACQUITY BEH C18 analytical column (1.7 μm particle size, 100 μm x 100 mm) over one hour with a gradient from 2% to 25% buffer B (99.9% ACN/0.1% formic acid) at a flow rate of 0.4 μl/min. Raw mass spectrometric data were converted into peak lists using Bioworks 3.3.1 SP1. The spectra were searched using Prospector v.5.3 (http://prospector.ucsf.edu) against a human-restricted UniProt database. Protein Prospector results were filtered by applying a minimum Protein Score of 22.0, a minimum Peptide Score of 15.0, a maximum Protein E-Value of 0.01 and a maximum Peptide E-Value of 0.05. A total of 419 AP-MS experiments were performed using each of the 90-tyrosine kinases as affinity-tagged bait. Preprocessing of IPs was carried out by mapping protein count obtained in individual replicate experiments corresponding to a bait. Based on the distribution of protein counts of all experiments we removed the samples where protein counts were less than 30. 19 AP-MS experiments fall under this category and they were removed from further analysis. Pull-down experiments where tyrosine kinase was not seen as bait in the mass spectra were discarded. Reproducibility of interactions of tyrosine kinases was defined as total number of preys picked by a kinase in at least two biological replicate experiments by the total number of preys picked by bait in experiments.

### Mass spectrometry data scoring

To distinguish true from non-specific interactions MS data was scored using two available scoring methods - Comparative Proteomics Analysis Software Suite (CompPASS) (Sowa et al., 2009) and Significance Analysis of Interactome (SAINT) (Choi et al., 2011). CompPASS scoring was performed as described by (Sowa et al. 2009). As the number of biological replicate experiments carried out for tyrosine kinases varied we modified the current CompPASS WD scores by adding the bait specific Z-normalization (CompPASS-Z). For every bait CompPASS WD scores were Z-scored to obtain modified CompPASS-Z scores as follows:

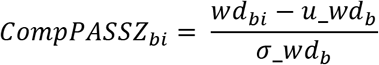

where *wd_bi_* represents the comPASS WD score for interaction between bait (*b*) and prey (*i*) and *wd_b_* represents all WD scores associated with bait *b*.

SAINT was also used to identify interactors from AP-MS data using SAINTexpress software (version 3, Teo et al). Performance of CompPASS, CompPASS-Z and SAINT scored AP-MS data was compared in scoring known interactions of HumanNet (Lee et al., 2011) and iRefWeb (Turner et al., 2010) databases as reference. The HumanNet v1 network was downloaded (10/27/2015) from http://www.functionalnet.org/humannet, iRefWeb v13.0 network was downloaded (7/23/2014) from http://www.irefindex.org, Bioplex was downloaded from http://bioplex.hms.harvard.edu/downloadInteractions.php (BioPlex_2.3_interactionList.tsv, 5/3/2018) (Huttlin et al., 2015, 2017), MINT database (Licata et al., 2012) was downloaded (8/11/2015) from https://mint.bio.uniroma2.it/ and had 1457 tyrosine kinase interactions. Statistical significance of overlap was obtained using hypergeometric test where background distribution was based on a random interaction network of uniquely identified preys in entire AP-MS with all 90 tyrosine kinases as baits. Similarly, we compared the overlap of CompPASS-Z scored AP-MS data with known kinase-substrate information downloaded from PhosphoSitePlus (Hornbeck et al., 2012) (downloaded on 8/5/2016, 699 tyrosine kinase substrates) at different CompPASS-Z cutoffs. A cutoff of CompPASS-Z scores was derived using a highest peak of enrichment using known interactors and substrates (Figure 1C). The number of high confidence interactors per kinase was not correlated with the number of replicates carried out per kinase (Pearson r = 0.05).

### Western blotting

Cell lysates were subjected to SDS-PAGE and transferred to nitrocellulose membranes. For signaling analysis by immunoblot in lung cancer cell lines, 2×10^6^ cells were seeded per 10cm dish for 24 hours after which cells were serum-starved for 18 hours and treated with either vehicle (DMSO) or Erlotinib (1μM) for one hour and stimulated with 100 ng/mL human recombinant EGF for five minutes prior to lysis.

### Cellular localization prediction

Protein subcellular localization were examined using the GO cellular components downloaded from Ensembl using R BioMart package (Durinck et al., 2005). To predict the cellular localization of a tyrosine kinase first degree (direct) interactors were used for enrichment analysis using Fisher’s Exact test. All the cellular localization terms were manually condensed to thirteen core localizations: Nucleus, Cytoplasm, Vesicles, Plasma membrane, ECM, Golgi, Endosomes, Cell junctions, Vacuoles, Cytoskeleton, Endoplasmic reticulum and exosomes. A tyrosine kinase was considered associated with a cellular localization when its interactors were significantly more likely to be annotated with the same cellular term as compared to human proteome (background). To focus on the most significant results, we used only cellular terms where enrichment scores were significant with an adjusted FDR of 10%.

### Network mapping of tyrosine kinases with protein complexes

To map the association of tyrosine kinases with known protein complexes we used known protein complex information from CORUM and iRefWeb databases. CORUM database (Giurgiu et al., 2019) was downloaded (3/18/2019) from http://mips.helmholtz-muenchen.de/corum/ with 6859 core complexes. Network representation was generated using Cytoscape version 3.2 (Shannon et al., 2003). Interaction coverage was defined as the number of interactors of a kinase which are a part of a protein complex over the total number of members of a protein complex. Statistical significance of the interaction overlap of a protein complexes was calculated using hypergeometric test and represents the enrichment of tyrosine kinase interactors to form a protein complex. All *P* values were FDR adjusted and a cutoff of 5% was used to identify tyrosine kinase-protein complex associations.

### Cell viability assay

Cells were seeded in 12-well format at 5,000 cells/well and treated with vehicle (DMSO) or erlotinib (1μM) on the following day. Media containing vehicle or drug, respectively, was replaced every 3-4 days and cells were maintained until full confluency of vehicle-treated control wells. Cells were fixed with 10% formalin for 30 minutes at room temperature and stained with 0.05% crystal violet solution. Pictures were taken using an ImageQuant LAS 4000 (GE Healthcare Life Sciences). For quantification, stained cells were dissolved with 1% SDS solution, and absorbance at 470 nm was measured using a Spectramax spectrophotometer (Molecular Devices). Values were normalized to vehicle-treated controls. For dacomitinib cell viability experiments a total of 2500-5000 cells were seeded per well, depending on their growth rates. After overnight incubation at 37°C, drugs were added to the culture wells to achieve a final drug concentration from 10 μM to 152 pM. After drug addition, the cells were incubated for an additional 3 days to allow cells to reach 70-90% confluency. Readout of cell viability was determined with 20 μL of 1 mg/mL resazurin salt (Sigma) using an Envision multi-reader. Cell counts were first adjusted by subtracting the average of the baseline cell counts from untreated cells assessed 1 day after cell seeding. A four-parameter logistic model was used to fit the dose response curves and infer the IC_50_, slope, and upper and lower limits.

### RNAi screening

Top high confidence interactors were selected for EGFR, ERBB2, ALK and MET tyrosine kinases based on the CompPASS-Z scores of the interactions. Cells were reverse transfected in quadruplicate in 384-well plates with 20nM of siRNA (Sigma) using RNAiMax as transfection reagent. Cells were transfected for 24 hr, and then the entire plate was treated with either one drug at a half maximal inhibitory concentration (IC_50_) concentration or DMSO for 72 hr, after which cells were stained with Hoescht 33342 and counted using a Thermo CellInsight high-content microscope. Both drug and DMSO plates for every cell line were median centered to normalize the proliferation rates. Next, four normalized replicate values in the DMSO plate were compared to the same gene in the drug plate and P value of the significance of this difference in medians were calculated using t-statistics. A *P* value of 0.05 was considered significant. The genetic interactions was defined by the negative log10 of the *P* value of t-test and signed with either positive (knockdown causes drug sensitivity) or negative (knockdown causes drug resistance).

### TCGA, cancer cell line and PDX datasets

Following datasets from TCGA and cancer cell lines (CCLE) were used to analyze the network activity of EGFR.

**Table 1.** TCGA datasets.

| **TCGA datasets** |  |  |
| --- | --- | --- |
| *Tumor type* | *Data type* | *# of samples* |
| LUAD | mRNA Expression | 575 |
|  | Mutation (EGFR) | 66 |
|  | Reverse Phase Protein Array (EGFR) | 360 |

**Table 2.** CCLE datasets.

| **CCLE datasets** |  |  |
| --- | --- | --- |
| *Tissue type* | *Data type* | *# of cell lines* |
| NSCLC | mRNA Expression | 51 |
|  | Mutation (EGFR) | 5 |
|  | Reverse Phase Protein Array (EGFR) | 51 |
|  | Drug response (Erlotinib/Lapatinib) | 29 |

LUAD patients mRNA, RPPA, CNV and mutation datasets were downloaded from TCGA data portal (https://portal.gdc.cancer.gov/). CCLE datasets were downloaded from (https://www.synapse.org/#!Synapse:syn10463688/wiki/463140). Gene expression datasets were log transformed and *z*-normalized across the NSCLC cell lines and patient samples. RPPA data of protein expression of EGFR in NSCLC cell lines was obtained from (https://bioinformatics.mdanderson.org/public-software/mclp/). PDX data was obtained from (Gao et al., 2015) and included baseline gene expression and tumor responses to erlotinib. Gene expression data was log transformed and normalized across all 25 NSCLC PDX models.

### Description of network activity scoring

To map the network activity of tyrosine kinases we developed a computational algorithm to integrate proteomics data (protein-protein interactors) of a tyrosine kinase with mRNA expression of tumor samples. This algorithm is based on the hypothesis that cellular activity of a kinase is dependent on its neighboring protein interactors and the levels of direct interactors of a kinase can influence the activity of the kinase itself. We refer to this interactome-based mapping of transcriptional levels as network activity and used this to classify tumors as network positive or negative.

To calculate network activity we first used mRNA expression of interactors of a tyrosine kinase in a specific tumor type. Here we focused on lung cancer where EGFR is a common mutational driver (~12% frequency). To identify the key interactors on which kinase activity is highly dependent, patients were separated into mutationally altered vs. non altered groups and mRNA expression of interactors was compared to identify interactors whose mean expression was significantly different between groups. Statistical significance of expression of interactors was achieved using a two-sided t-test with a FDR cutoff of 5%. Positive component (*pc*) comprises interactors whose expression was elevated when the tyrosine kinase is altered, while negative component (*nc*) includes interactors whose expression was reduced when the kinase was altered (i.e higher in wild type samples). To generate the network activity score for sample *k* (*NA_k_*) we sum up the expression of interactors of positive and negative components and calculate the difference in combined expression of two components as follows:

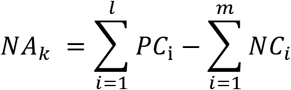

where, *PC* and *NC* represent the expression of interactors in *pc* and *nc* respectively. *l* and *m* represents the number of interactors in positive and negative components respectively.

Interactors of EGFR obtained through our AP-MS study at CompPASS-Z cutoff of 2 and other experimentally known interactors of EGFR from iRefWeb database (total n = 181) were used to calculate network activity. Out of 181 direct interactors of EGFR, 27 interactors passed the FDR threshold of 5% and were used for NA calculations of EGFR in LUAD. A cutoff for NA score was calculated by precision-recall analysis by considering canonical mutations of EGFR as true positives and KRAS mutations as false positives. A recall of 95% was considered to derive a NA cutoff in LUAD patients

### Comparison of EGFR network activity with other methods

Performance of EGFR NA was compared to either random gene networks or other gene expression signature approaches. To compare to random gene networks we selected 27 genes at random whose expression was differentially expressed in samples with an EGFR mutation, as above. This process was performed 10,000 times and the predictive capacity of each trial was compared to the real EGFR NA. The percent of random trials where the prediction was stronger than the real EGFR NA was used to calculate a *P* value.

To compare to other gene expression signatures approaches we first identified which of all 20,000 genes were differentially expressed genes (n =3299) between EGFR wild-type and mutant samples at a 5% FDR threshold. The gene expression signature (GES) was calculated by a signed sum of differentially expressed genes. LASSO, ElasticNet and Random Forest classifier models were generated by applying these methods to the 20,000 gene set trained on EGFR mutant versus wild-type samples. LASSO was run with ∝ =0.01 and maximum iterations = 10e5 where ∝ was chosen by gridsearch during training. ElasticNet was run with l1_ratio =0.5 and ∝ =0.01, which was chosen based on internal cross validation. Random Forest model was built with 100 estimators and gini criterion. For Random Forest average expression of the most predictive genes was used to calculate EGFR gene expression signature. EGFR activity using these methods was compared to pEGFR and drug response data from CCLE and PDX models.

### NMF and mutation enrichment analysis of EGFR^wt,n+^ patients

Mutations enrichment in patients with high activity of EGFR was calculated by measuring the fraction of mutations in samples with high NA as compared to background (total mutant samples of a gene) mutation frequency of a gene in all LUAD samples. Significance of the mutation enrichment was calculated using Hypergeometric test. Mutation enrichment analysis for genes except EGFR was performed in EGFR null samples. Similarly, mutation frequency of specific alleles of KRAS and BRAF was calculated in EGFR^wt,n+^ patients. Background mutation frequency represents total KRAS and BRAF mutation frequency in patients with high EGFR NA over background. EGFR NA high patient samples (n=148) were decomposed using Non-Negative Matrix Factorization (NMF) (Brunet et al., 2004; Kim et al., 2017; Tamayo et al., 2007). Matrix (A) contains *n* rows (expression of EGFR interactors which are part of NA, n=27) and *m* samples (EGFR mutant and EGFR^wt,n+^ samples, n=148) and is represented as A^nXM^. NMF was implemented as in (Kim et al., 2017). The optimal number of components was defined using cophenetic correlation (Brunet et al. 2004). Mutation enrichment analysis was performed in every state to identify correspondence between the present of a somatic mutation and EGFR network state. Enrichment analysis was performed in every state by considering other states as background and genes which are significantly mutated were identified using Fischer’s Exact test (p <0.01).

### Mouse xenograft study

Six- to seven-week-old female nude mice were purchased from the Jackson Laboratory and housed with ad libitum food and water on a 12-hour light cycle at the UCSF Preclinical Therapeutics Core vivarium. All animal studies were performed in full accordance with UCSF Institutional Animal Care and Use Committee. H358 xenografts were established by subcutaneous injection into the left and right flanks of mice with H358 cells (5 × 10^6^ cells in 100 μl of serum-free medium mixed 1:1 with Matrigel). Tumor xenografts were allowed to establish until they reached about 700 to 900 mm^3^ in donor mice and then reimplanted into receiver mice to achieve higher engraftment rate. Briefly, established H538 tumor xenografts from donor mice were resected, cut into even-size fragments (15 mm × 15 mm), embedded in Matrigel, and reimplanted via subcutaneous implantation into receiver mice. H538 tumor-bearing mice were randomized into control and treatment groups when tumors reached a size range of 100 to 120 mm^3^, and single or dual dosing of erlotinib (80-100 mg/kg in 0.5% (hydroxypropyl) methyl cellulose, 0.2% Tween 80 (HPMT)] or vehicle control (Labrasol) was administered daily by oral gavage. Tumor volume and body weight were assessed biweekly for the duration of the study. The percentage change in tumor growth was based on volumes calculated from the size on day 1 at the beginning of treatment. For PDX studies, patient-derived tumor cells were engrafted subcutaneously into the flank of C.B-17 SCID mice. Tumors were allowed to grow until they reached a minimum volume of 200 mm^3^; then, animals were randomly placed into control or treatment groups. Animals were treated daily for 30 d via oral gavage, and tumor volume was calculated daily using caliper measurements.

### Data availability

Network data is deposited in Biogrid (Accession Code XXXXX). Source code for Compass-Z, Non-negative matrix factorization and Network Activity analysis are available at https://github.com/BandyopadhyayLab/Tyrosine_kinase_interactome

## REFERENCES

1. Bache, K.G., Slagsvold, T., and Stenmark, H. (2004). Defective downregulation of receptor tyrosine kinases in cancer. EMBO J. 23, 2707–2712.

2. Bajenaru, M.L., Zhu, Y., Hedrick, N.M., Donahoe, J., Parada, L.F., and Gutmann, D.H. (2002). Astrocyte-specific inactivation of the neurofibromatosis 1 gene (NF1) is insufficient for astrocytoma formation. Mol. Cell. Biol. 22, 5100–5113.

3. Bandyopadhyay, S., Chiang, C., Srivastava, J., Gersten, M., White, S., Bell, R., Kurschner, C., Martin, C.H., Smoot, M., Sahasrabudhe, S., et al. (2010). A human MAP kinase interactome. Nat. Methods 7, 801–805.

4. Barretina, J., Caponigro, G., Stransky, N., Venkatesan, K., Margolin, A.A., Kim, S., Wilson, C.J., Lehár, J., Kryukov, G.V., Sonkin, D., et al. (2012). The Cancer Cell Line Encyclopedia enables predictive modelling of anticancer drug sensitivity. Nature 483, 603–607.

5. Bartholomeeusen, K., Xiang, Y., Fujinaga, K., and Peterlin, B.M. (2012). Bromodomain and extra-terminal (BET) bromodomain inhibition activate transcription via transient release of positive transcription elongation factor b (P-TEFb) from 7SK small nuclear ribonucleoprotein. J. Biol. Chem. 287, 36609–36616.

6. Batzer, A.G., Rotin, D., Ureña, J.M., Skolnik, E.Y., and Schlessinger, J. (1994). Hierarchy of binding sites for Grb2 and Shc on the epidermal growth factor receptor. Mol. Cell. Biol. 14, 5192–5201.

7. Behrends, C., Sowa, M.E., Gygi, S.P., and Harper, J.W. (2010). Network organization of the human autophagy system. Nature 466, 68–76.

8. Bello, C.L., Smith, E., Ruiz-Garcia, A., Ni, G., Alvey, C., and Loi, C.-M. (2013). A phase I, open-label, mass balance study of [(14)C] dacomitinib (PF-00299804) in healthy male volunteers. Cancer Chemother. Pharmacol. 72, 379–385.

9. Berger, A.H., Brooks, A.N., Wu, X., Shrestha, Y., Chouinard, C., Piccioni, F., Bagul, M., Kamburov, A., Imielinski, M., Hogstrom, L., et al. (2016). High-throughput Phenotyping of Lung Cancer Somatic Mutations. Cancer Cell 30, 214–228.

10. Bild, A.H., Yao, G., Chang, J.T., Wang, Q., Potti, A., Chasse, D., Joshi, M.-B., Harpole, D., Lancaster, J.M., Berchuck, A., et al. (2006). Oncogenic pathway signatures in human cancers as a guide to targeted therapies. Nature 439, 353–357.

11. Bollag, G., Clapp, D.W., Shih, S., Adler, F., Zhang, Y.Y., Thompson, P., Lange, B.J., Freedman, M.H., McCormick, F., Jacks, T., et al. (1996). Loss of NF1 results in activation of the Ras signaling pathway and leads to aberrant growth in haematopoietic cells. Nat. Genet. 12, 144–148.

12. Breitkreutz, A., Choi, H., Sharom, J.R., Boucher, L., Neduva, V., Larsen, B., Lin, Z.-Y., Breitkreutz, B.-J., Stark, C., Liu, G., et al. (2010). A global protein kinase and phosphatase interaction network in yeast. Science 328, 1043–1046.

13. Cancer Genome Atlas Research Network (2014). Comprehensive molecular profiling of lung adenocarcinoma. Nature 511, 543–550.

14. Carpenter, G. (2003). Nuclear localization and possible functions of receptor tyrosine kinases. Curr. Opin. Cell Biol. 15, 143–148.

15. Carpenter, G., and Liao, H.-J. (2013). Receptor tyrosine kinases in the nucleus. Cold Spring Harb. Perspect. Biol. 5, a008979.

16. Du, Z., and Lovly, C.M. (2018). Mechanisms of receptor tyrosine kinase activation in cancer. Mol. Cancer 17, 58.

17. Ferté, C., Besse, B., Dansin, E., Parent, F., Buisine, M.-P., Copin, M.-C., Penel, N., and Soria, J.-C. (2010). Durable responses to Erlotinib despite KRAS mutations in two patients with metastatic lung adenocarcinoma. Ann. Oncol. Off. J. Eur. Soc. Med. Oncol. 21, 1385–1387.

18. Fujimoto, N., Wislez, M., Zhang, J., Iwanaga, K., Dackor, J., Hanna, A.E., Kalyankrishna, S., Cody, D.D., Price, R.E., Sato, M., et al. (2005). High Expression of ErbB Family Members and Their Ligands in Lung Adenocarcinomas That Are Sensitive to Inhibition of Epidermal Growth Factor Receptor. Cancer Res. 65, 11478–11485.

19. Gao, H., Korn, J.M., Ferretti, S., Monahan, J.E., Wang, Y., Singh, M., Zhang, C., Schnell, C., Yang, G., Zhang, Y., et al. (2015). High-throughput screening using patient-derived tumor xenografts to predict clinical trial drug response. Nat. Med. 21, 1318–1325.

20. Gschwind, A., Fischer, O.M., and Ullrich, A. (2004). The discovery of receptor tyrosine kinases: targets for cancer therapy. Nat. Rev. Cancer 4, 361–370.

21. Haigis, K.M. (2017). KRAS Alleles: The Devil Is in the Detail. Trends Cancer 3, 686–697.

22. Hein, M.Y., Hubner, N.C., Poser, I., Cox, J., Nagaraj, N., Toyoda, Y., Gak, I.A., Weisswange, I., Mansfeld, J., Buchholz, F., et al. (2015). A human interactome in three quantitative dimensions organized by stoichiometries and abundances. Cell 163, 712–723.

23. Hirai, F., Edagawa, M., Shimamatsu, S., Toyozawa, R., Toyokawa, G., Nosaki, K., Yamaguchi, M., Seto, T., Takenoyama, M., and Ichinose, Y. (2017). Evaluation of erlotinib for the treatment of patients with non-small cell lung cancer with epidermal growth factor receptor wild type. Oncol. Lett. 14, 306–312.

24. Hornbeck, P.V., Kornhauser, J.M., Tkachev, S., Zhang, B., Skrzypek, E., Murray, B., Latham, V., and Sullivan, M. (2012). PhosphoSitePlus: a comprehensive resource for investigating the structure and function of experimentally determined post-translational modifications in man and mouse. Nucleic Acids Res. 40, D261–270.

25. van Houdt, W.J., Hoogwater, F.J.H., de Bruijn, M.T., Emmink, B.L., Nijkamp, M.W., Raats, D.A.E., van der Groep, P., van Diest, P., Borel Rinkes, I.H.M., and Kranenburg, O. (2010). Oncogenic KRAS desensitizes colorectal tumor cells to epidermal growth factor receptor inhibition and activation. Neoplasia N. Y. N 12, 443–452.

26. Hu, H.-M., Zhao, X., Kaushik, S., Robillard, L., Barthelet, A., Lin, K.K., Shah, K.N., Simmons, A.D., Raponi, M., Harding, T.C., et al. (2018). A Quantitative Chemotherapy Genetic Interaction Map Reveals Factors Associated with PARP Inhibitor Resistance. Cell Rep. 23, 918–929.

27. Hunter, J.C., Manandhar, A., Carrasco, M.A., Gurbani, D., Gondi, S., and Westover, K.D. (2015). Biochemical and Structural Analysis of Common Cancer-Associated KRAS Mutations. Mol. Cancer Res. MCR 13, 1325–1335.

28. Huttlin, E.L., Ting, L., Bruckner, R.J., Gebreab, F., Gygi, M.P., Szpyt, J., Tam, S., Zarraga, G., Colby, G., Baltier, K., et al. (2015). The BioPlex Network: A Systematic Exploration of the Human Interactome. Cell 162, 425–440.

29. Huttlin, E.L., Bruckner, R.J., Paulo, J.A., Cannon, J.R., Ting, L., Baltier, K., Colby, G., Gebreab, F., Gygi, M.P., Parzen, H., et al. (2017). Architecture of the human interactome defines protein communities and disease networks. Nature 545, 505–509.

30. James, R.G., Biechele, T.L., Conrad, W.H., Camp, N.D., Fass, D.M., Major, M.B., Sommer, K., Yi, X., Roberts, B.S., Cleary, M.A., et al. (2009). Bruton’s tyrosine kinase revealed as a negative regulator of Wnt-beta-catenin signaling. Sci. Signal. 2, ra25.

31. Janes, M.R., Zhang, J., Li, L.-S., Hansen, R., Peters, U., Guo, X., Chen, Y., Babbar, A., Firdaus, S.J., Darjania, L., et al. (2018). Targeting KRAS Mutant Cancers with a Covalent G12C-Specific Inhibitor. Cell 172, 578–589.e17.

32. Jazieh, A.-R., Al Sudairy, R., Abu-Shraie, N., Al Suwairi, W., Ferwana, M., and Murad, M.H. (2013). Erlotinib in wild type epidermal growth factor receptor non-small cell lung cancer: A systematic review. Ann. Thorac. Med. 8, 204–208.

33. Kim, J.W., Abudayyeh, O.O., Yeerna, H., Yeang, C.-H., Stewart, M., Jenkins, R.W., Kitajima, S., Konieczkowski, D.J., Medetgul-Ernar, K., Cavazos, T., et al. (2017). Decomposing Oncogenic Transcriptional Signatures to Generate Maps of Divergent Cellular States. Cell Syst. 5, 105–118.e9.

34. Koboldt, D.C., Fulton, R.S., McLellan, M.D., Schmidt, H., Kalicki-Veizer, J., McMichael, J.F., Fulton, L.L., Dooling, D.J., Ding, L., Mardis, E.R., et al. (2012). Comprehensive molecular portraits of human breast tumours. Nature 490, 61–70.

35. Kohsaka, S., Nagano, M., Ueno, T., Suehara, Y., Hayashi, T., Shimada, N., Takahashi, K., Suzuki, K., Takamochi, K., Takahashi, F., et al. (2017). A method of high-throughput functional evaluation of EGFR gene variants of unknown significance in cancer. Sci. Transl. Med. 9.

36. Krejci, J., Pesek, M., Grossmann, P., Krejci, M., Ricar, J., Benesova, L., and Minarik, M. (2011). Extraordinary response to erlotinib therapy in a patient with lung adenocarcinoma exhibiting KRAS mutation and EGFR amplification. Cancer Genomics Proteomics 8, 135–138.

37. Krogan, N.J., Cagney, G., Yu, H., Zhong, G., Guo, X., Ignatchenko, A., Li, J., Pu, S., Datta, N., Tikuisis, A.P., et al. (2006). Global landscape of protein complexes in the yeast Saccharomyces cerevisiae. Nature 440, 637–643.

38. Kruspig, B., Monteverde, T., Neidler, S., Hock, A., Kerr, E., Nixon, C., Clark, W., Hedley, A., Laing, S., Coffelt, S.B., et al. (2018). The ERBB network facilitates KRAS-driven lung tumorigenesis. Sci. Transl. Med. 10.

39. Largaespada, D.A., Brannan, C.I., Jenkins, N.A., and Copeland, N.G. (1996). Nf1 deficiency causes Ras-mediated granulocyte/macrophage colony stimulating factor hypersensitivity and chronic myeloid leukaemia. Nat. Genet. 12, 137–143.

40. Lee, I., Blom, U.M., Wang, P.I., Shim, J.E., and Marcotte, E.M. (2011). Prioritizing candidate disease genes by network-based boosting of genome-wide association data. Genome Res. 21, 1109–1121.

41. Li, X., Huang, Y., Jiang, J., and Frank, S.J. (2008). ERK-dependent threonine phosphorylation of EGF receptor modulates receptor downregulation and signaling. Cell. Signal. 20, 2145–2155.

42. Loboda, A., Nebozhyn, M., Klinghoffer, R., Frazier, J., Chastain, M., Arthur, W., Roberts, B., Zhang, T., Chenard, M., Haines, B., et al. (2010). A gene expression signature of RAS pathway dependence predicts response to PI3K and RAS pathway inhibitors and expands the population of RAS pathway activated tumors. BMC Med. Genomics 3, 26.

43. Lou, K., Steri, V., Ge, A.Y., Hwang, Y.C., Yogodzinski, C.H., Shkedi, A.R., Choi, A.L.M., Mitchell, D.C., Swaney, D.L., Hann, B., et al. (2019). KRASG12C inhibition produces a driver-limited state revealing collateral dependencies. Sci. Signal. 12.

44. Lynch, T.J., Bell, D.W., Sordella, R., Gurubhagavatula, S., Okimoto, R.A., Brannigan, B.W., Harris, P.L., Haserlat, S.M., Supko, J.G., Haluska, F.G., et al. (2004). Activating mutations in the epidermal growth factor receptor underlying responsiveness of non-small-cell lung cancer to gefitinib. N. Engl. J. Med. 350, 2129–2139.

45. Marquart, J., Chen, E.Y., and Prasad, V. (2018). Estimation of the Percentage of US Patients With Cancer Who Benefit From Genome-Driven Oncology. JAMA Oncol. 4, 1093–1098.

46. Marshall, N.F., Peng, J., Xie, Z., and Price, D.H. (1996). Control of RNA polymerase II elongation potential by a novel carboxyl-terminal domain kinase. J. Biol. Chem. 271, 27176–27183.

47. Mirza, M.R., Monk, B.J., Herrstedt, J., Oza, A.M., Mahner, S., Redondo, A., Fabbro, M., Ledermann, J.A., Lorusso, D., Vergote, I., et al. (2016). Niraparib Maintenance Therapy in Platinum-Sensitive, Recurrent Ovarian Cancer. N. Engl. J. Med. 375, 2154–2164.

48. Moll, H.P., Pranz, K., Musteanu, M., Grabner, B., Hruschka, N., Mohrherr, J., Aigner, P., Stiedl, P., Brcic, L., Laszlo, V., et al. (2018). Afatinib restrains K-RAS-driven lung tumorigenesis. Sci. Transl. Med. 10.

49. Moroni, M., Veronese, S., Benvenuti, S., Marrapese, G., Sartore-Bianchi, A., Di Nicolantonio, F., Gambacorta, M., Siena, S., and Bardelli, A. (2005). Gene copy number for epidermal growth factor receptor (EGFR) and clinical response to antiEGFR treatment in colorectal cancer: a cohort study. Lancet Oncol. 6, 279–286.

50. Ng, P.K.-S., Li, J., Jeong, K.J., Shao, S., Chen, H., Tsang, Y.H., Sengupta, S., Wang, Z., Bhavana, V.H., Tran, R., et al. (2018). Systematic Functional Annotation of Somatic Mutations in Cancer. Cancer Cell 33, 450–462.e10.

51. Ni, C.Y., Murphy, M.P., Golde, T.E., and Carpenter, G. (2001). gamma -Secretase cleavage and nuclear localization of ErbB-4 receptor tyrosine kinase. Science 294, 2179–2181.

52. Nichols, R.J., Haderk, F., Stahlhut, C., Schulze, C.J., Hemmati, G., Wildes, D., Tzitzilonis, C., Mordec, K., Marquez, A., Romero, J., et al. (2018). RAS nucleotide cycling underlies the SHP2 phosphatase dependence of mutant BRAF-, NF1- and RAS-driven cancers. Nat. Cell Biol. 20, 1064–1073.

53. Osarogiagbon, R.U., Cappuzzo, F., Ciuleanu, T., Leon, L., and Klughammer, B. (2015). Erlotinib therapy after initial platinum doublet therapy in patients with EGFR wild type non-small cell lung cancer: results of a combined patient-level analysis of the NCIC CTG BR.21 and SATURN trials. Transl. Lung Cancer Res. 4, 465–474.

54. Pao, W., Miller, V., Zakowski, M., Doherty, J., Politi, K., Sarkaria, I., Singh, B., Heelan, R., Rusch, V., Fulton, L., et al. (2004). EGF receptor gene mutations are common in lung cancers from “never smokers” and are associated with sensitivity of tumors to gefitinib and erlotinib. Proc. Natl. Acad. Sci. U. S. A. 101, 13306–13311.

55. Paul, M.D., and Hristova, K. (2019). The RTK Interactome: Overview and Perspective on RTK Heterointeractions. Chem. Rev. 119, 5881–5921.

56. Popovici, V., Budinska, E., Tejpar, S., Weinrich, S., Estrella, H., Hodgson, G., Van Cutsem, E., Xie, T., Bosman, F.T., Roth, A.D., et al. (2012). Identification of a poor-prognosis BRAF-mutant-like population of patients with colon cancer. J. Clin. Oncol. Off. J. Am. Soc. Clin. Oncol. 30, 1288–1295.

57. Robinson, D.R., Wu, Y.M., and Lin, S.F. (2000). The protein tyrosine kinase family of the human genome. Oncogene 19, 5548–5557.

58. Rojas, M., Yao, S., and Lin, Y.Z. (1996). Controlling epidermal growth factor (EGF)-stimulated Ras activation in intact cells by a cell-permeable peptide mimicking phosphorylated EGF receptor. J. Biol. Chem. 271, 27456–27461.

59. Rual, J.-F., Venkatesan, K., Hao, T., Hirozane-Kishikawa, T., Dricot, A., Li, N., Berriz, G.F., Gibbons, F.D., Dreze, M., Ayivi-Guedehoussou, N., et al. (2005). Towards a proteome-scale map of the human protein-protein interaction network. Nature 437, 1173–1178.

60. Sanchez-Vega, F., Mina, M., Armenia, J., Chatila, W.K., Luna, A., La, K.C., Dimitriadoy, S., Liu, D.L., Kantheti, H.S., Saghafinia, S., et al. (2018). Oncogenic Signaling Pathways in The Cancer Genome Atlas. Cell 173, 321–337.e10.

61. Schwikowski, B., Uetz, P., and Fields, S. (2000). A network of protein-protein interactions in yeast. Nat. Biotechnol. 18, 1257–1261.

62. Shah, S.P., Roth, A., Goya, R., Oloumi, A., Ha, G., Zhao, Y., Turashvili, G., Ding, J., Tse, K., Haffari, G., et al. (2012). The clonal and mutational evolution spectrum of primary triple-negative breast cancers. Nature 486, 395–399.

63. Shao, Y.-Y., Hsu, C.-H., and Cheng, A.-L. (2015). Predictive biomarkers of sorafenib efficacy in advanced hepatocellular carcinoma: Are we getting there? World J. Gastroenterol. WJG 21, 10336–10347.

64. Shin, C.J., Wong, S., Davis, M.J., and Ragan, M.A. (2009). Protein-protein interaction as a predictor of subcellular location. BMC Syst. Biol. 3, 28.

65. Singh, A.B., and Harris, R.C. (2005). Autocrine, paracrine and juxtacrine signaling by EGFR ligands. Cell. Signal. 17, 1183–1193.

66. Sowa, M.E., Bennett, E.J., Gygi, S.P., and Harper, J.W. (2009). Defining the human deubiquitinating enzyme interaction landscape. Cell 138, 389–403.

67. Stuart, J.R., Gonzalez, F.H., Kawai, H., and Yuan, Z.-M. (2006). c-Abl interacts with the WAVE2 signaling complex to induce membrane ruffling and cell spreading. J. Biol. Chem. 281, 31290–31297.

68. Sunaga, N., Shames, D.S., Girard, L., Peyton, M., Larsen, J.E., Imai, H., Soh, J., Sato, M., Yanagitani, N., Kaira, K., et al. (2011). Knockdown of oncogenic KRAS in non-small cell lung cancers suppresses tumor growth and sensitizes tumor cells to targeted therapy. Mol. Cancer Ther. 10, 336–346.

69. Swisher, E.M., Lin, K.K., Oza, A.M., Scott, C.L., Giordano, H., Sun, J., Konecny, G.E., Coleman, R.L., Tinker, A.V., O’Malley, D.M., et al. (2017). Rucaparib in relapsed, platinum-sensitive high-grade ovarian carcinoma (ARIEL2 Part 1): an international, multicentre, open-label, phase 2 trial. Lancet Oncol. 18, 75–87.

70. Tanos, B., and Pendergast, A.M. (2006). Abl tyrosine kinase regulates endocytosis of the epidermal growth factor receptor. J. Biol. Chem. 281, 32714–32723.

71. Thul, P.J., Åkesson, L., Wiking, M., Mahdessian, D., Geladaki, A., Ait Blal, H., Alm, T., Asplund, A., Björk, L., Breckels, L.M., et al. (2017). A subcellular map of the human proteome. Science 356.

72. Tian, S., Simon, I., Moreno, V., Roepman, P., Tabernero, J., Snel, M., van’t Veer, L., Salazar, R., Bernards, R., and Capella, G. (2013). A combined oncogenic pathway signature of BRAF, KRAS and PI3KCA mutation improves colorectal cancer classification and cetuximab treatment prediction. Gut 62, 540–549.

73. Turner, B., Razick, S., Turinsky, A.L., Vlasblom, J., Crowdy, E.K., Cho, E., Morrison, K., Donaldson, I.M., and Wodak, S.J. (2010). iRefWeb: interactive analysis of consolidated protein interaction data and their supporting evidence. Database J. Biol. Databases Curation 2010, baq023.

74. Wu, W., O’Reilly, M.S., Langley, R.R., Tsan, R.Z., Baker, C.H., Bekele, N., Tang, X.M., Onn, A., Fidler, I.J., and Herbst, R.S. (2007). Expression of epidermal growth factor (EGF)/transforming growth factor-α by human lung cancer cells determines their response to EGF receptor tyrosine kinase inhibition in the lungs of mice. Mol. Cancer Ther. 6, 2652–2663.

75. Yaffe, M.B. (2013). The scientific drunk and the lamppost: massive sequencing efforts in cancer discovery and treatment. Sci. Signal. 6, pe13.

76. Yao, Z., Darowski, K., St-Denis, N., Wong, V., Offensperger, F., Villedieu, A., Amin, S., Malty, R., Aoki, H., Guo, H., et al. (2017a). A Global Analysis of the Receptor Tyrosine Kinase-Protein Phosphatase Interactome. Mol. Cell 65, 347–360.

77. Yao, Z., Yaeger, R., Rodrik-Outmezguine, V.S., Tao, A., Torres, N.M., Chang, M.T., Drosten, M., Zhao, H., Cecchi, F., Hembrough, T., et al. (2017b). Tumours with class 3 BRAF mutants are sensitive to the inhibition of activated RAS. Nature 548, 234–238.

78. Zhang, G., Scarborough, H., Kim, J., Rozhok, A.I., Chen, Y.A., Zhang, X., Song, L., Bai, Y., Fang, B., Liu, R.Z., et al. (2016). Coupling an EML4-ALK-centric interactome with RNA interference identifies sensitizers to ALK inhibitors. Sci. Signal. 9, rs12.

## METHODS REFERENCES

1. Brunet, J.-P., Tamayo, P., Golub, T.R., and Mesirov, J.P. (2004). Metagenes and molecular pattern discovery using matrix factorization. Proc. Natl. Acad. Sci. U. S. A. 101, 4164–4169.

2. Choi, H., Larsen, B., Lin, Z.-Y., Breitkreutz, A., Mellacheruvu, D., Fermin, D., Qin, Z.S., Tyers, M., Gingras, A.-C., and Nesvizhskii, A.I. (2011). SAINT: probabilistic scoring of affinity purification-mass spectrometry data. Nat. Methods 8, 70–73.

3. Cichocki, A., Zdunek, R., and Amari, S. (2008). Nonnegative Matrix and Tensor Factorization [Lecture Notes]. IEEE Signal Process. Mag. 25, 142–145.

4. Durinck, S., Moreau, Y., Kasprzyk, A., Davis, S., De Moor, B., Brazma, A., and Huber, W. (2005). BioMart and Bioconductor: a powerful link between biological databases and microarray data analysis. Bioinforma. Oxf. Engl. 21, 3439–3440.

5. Gao, H., Korn, J.M., Ferretti, S., Monahan, J.E., Wang, Y., Singh, M., Zhang, C., Schnell, C., Yang, G., Zhang, Y., et al. (2015). High-throughput screening using patient-derived tumor xenografts to predict clinical trial drug response. Nat. Med. 21, 1318–1325.

6. Giurgiu, M., Reinhard, J., Brauner, B., Dunger-Kaltenbach, I., Fobo, G., Frishman, G., Montrone, C., and Ruepp, A. (2019). CORUM: the comprehensive resource of mammalian protein complexes-2019. Nucleic Acids Res. 47, D559–D563.

7. Hein, M.Y., Hubner, N.C., Poser, I., Cox, J., Nagaraj, N., Toyoda, Y., Gak, I.A., Weisswange, I., Mansfeld, J., Buchholz, F., et al. (2015). A human interactome in three quantitative dimensions organized by stoichiometries and abundances. Cell 163, 712–723.

8. Hornbeck, P.V., Kornhauser, J.M., Tkachev, S., Zhang, B., Skrzypek, E., Murray, B., Latham, V., and Sullivan, M. (2012). PhosphoSitePlus: a comprehensive resource for investigating the structure and function of experimentally determined post-translational modifications in man and mouse. Nucleic Acids Res. 40, D261–270.

9. Huttlin, E.L., Ting, L., Bruckner, R.J., Gebreab, F., Gygi, M.P., Szpyt, J., Tam, S., Zarraga, G., Colby, G., Baltier, K., et al. (2015). The BioPlex Network: A Systematic Exploration of the Human Interactome. Cell 162, 425–440.

10. Huttlin, E.L., Bruckner, R.J., Paulo, J.A., Cannon, J.R., Ting, L., Baltier, K., Colby, G., Gebreab, F., Gygi, M.P., Parzen, H., et al. (2017). Architecture of the human interactome defines protein communities and disease networks. Nature 545, 505–509.

11. Kim, J.W., Abudayyeh, O.O., Yeerna, H., Yeang, C.-H., Stewart, M., Jenkins, R.W., Kitajima, S., Konieczkowski, D.J., Medetgul-Ernar, K., Cavazos, T., et al. (2017). Decomposing Oncogenic Transcriptional Signatures to Generate Maps of Divergent Cellular States. Cell Syst. 5, 105–118.e9.

12. Lee, I., Blom, U.M., Wang, P.I., Shim, J.E., and Marcotte, E.M. (2011). Prioritizing candidate disease genes by network-based boosting of genome-wide association data. Genome Res. 21, 1109–1121.

13. Licata, L., Briganti, L., Peluso, D., Perfetto, L., Iannuccelli, M., Galeota, E., Sacco, F., Palma, A., Nardozza, A.P., Santonico, E., et al. (2012). MINT, the molecular interaction database: 2012 update. Nucleic Acids Res. 40, D857–D861.

14. Rual, J.-F., Venkatesan, K., Hao, T., Hirozane-Kishikawa, T., Dricot, A., Li, N., Berriz, G.F., Gibbons, F.D., Dreze, M., Ayivi-Guedehoussou, N., et al. (2005). Towards a proteome-scale map of the human protein-protein interaction network. Nature 437, 1173–1178.

15. Shannon, P., Markiel, A., Ozier, O., Baliga, N.S., Wang, J.T., Ramage, D., Amin, N., Schwikowski, B., and Ideker, T. (2003). Cytoscape: A Software Environment for Integrated Models of Biomolecular Interaction Networks. Genome Res. 13, 2498–2504.

16. Sowa, M.E., Bennett, E.J., Gygi, S.P., and Harper, J.W. (2009). Defining the human deubiquitinating enzyme interaction landscape. Cell 138, 389–403.

17. Tamayo, P., Scanfeld, D., Ebert, B.L., Gillette, M.A., Roberts, C.W.M., and Mesirov, J.P. (2007). Metagene projection for cross-platform, cross-species characterization of global transcriptional states. Proc. Natl. Acad. Sci. 104, 5959–5964.

18. Turner, B., Razick, S., Turinsky, A.L., Vlasblom, J., Crowdy, E.K., Cho, E., Morrison, K., Donaldson, I.M., and Wodak, S.J. (2010). iRefWeb: interactive analysis of consolidated protein interaction data and their supporting evidence. Database J. Biol. Databases Curation 2010, baq023.

19. Uhlén, M., Björling, E., Agaton, C., Szigyarto, C.A.-K., Amini, B., Andersen, E., Andersson, A.-C., Angelidou, P., Asplund, A., Asplund, C., et al. (2005). A Human Protein Atlas for Normal and Cancer Tissues Based on Antibody Proteomics. Mol. Cell. Proteomics 4, 1920–1932.

20. Uhlén, M., Fagerberg, L., Hallström, B.M., Lindskog, C., Oksvold, P., Mardinoglu, A., Sivertsson, Å., Kampf, C., Sjöstedt, E., Asplund, A., et al. (2015). Tissue-based map of the human proteome. Science 347.

