## Supplementary Figures for "A tyrosine kinase protein interaction map reveals targetable EGFR network oncogenesis in lung cancer"

### SUPPLEMENTARY TABLES

**Supplementary Table 1:** List of tyrosine kinase interactors identified in TyKiNet.

**Supplementary Table 2:** CORUM complexes that interact with tyrosine kinases.

**Supplementary Table 3:** EGFR network scores in TCGA LUAD samples.

**Supplementary Table 4:** EGFR network activity scores in NSCLC PDX models.

**Supplementary Table 5:** State assignments and mutational analysis in $EGFR^{wt,n+}$ LUAD patients.

**Supplementary Table 6:** Fitted dacomitinib drug sensitivity data in KRAS mutant NSCLC cell lines.

### SUPPLEMENTARY FIGURES

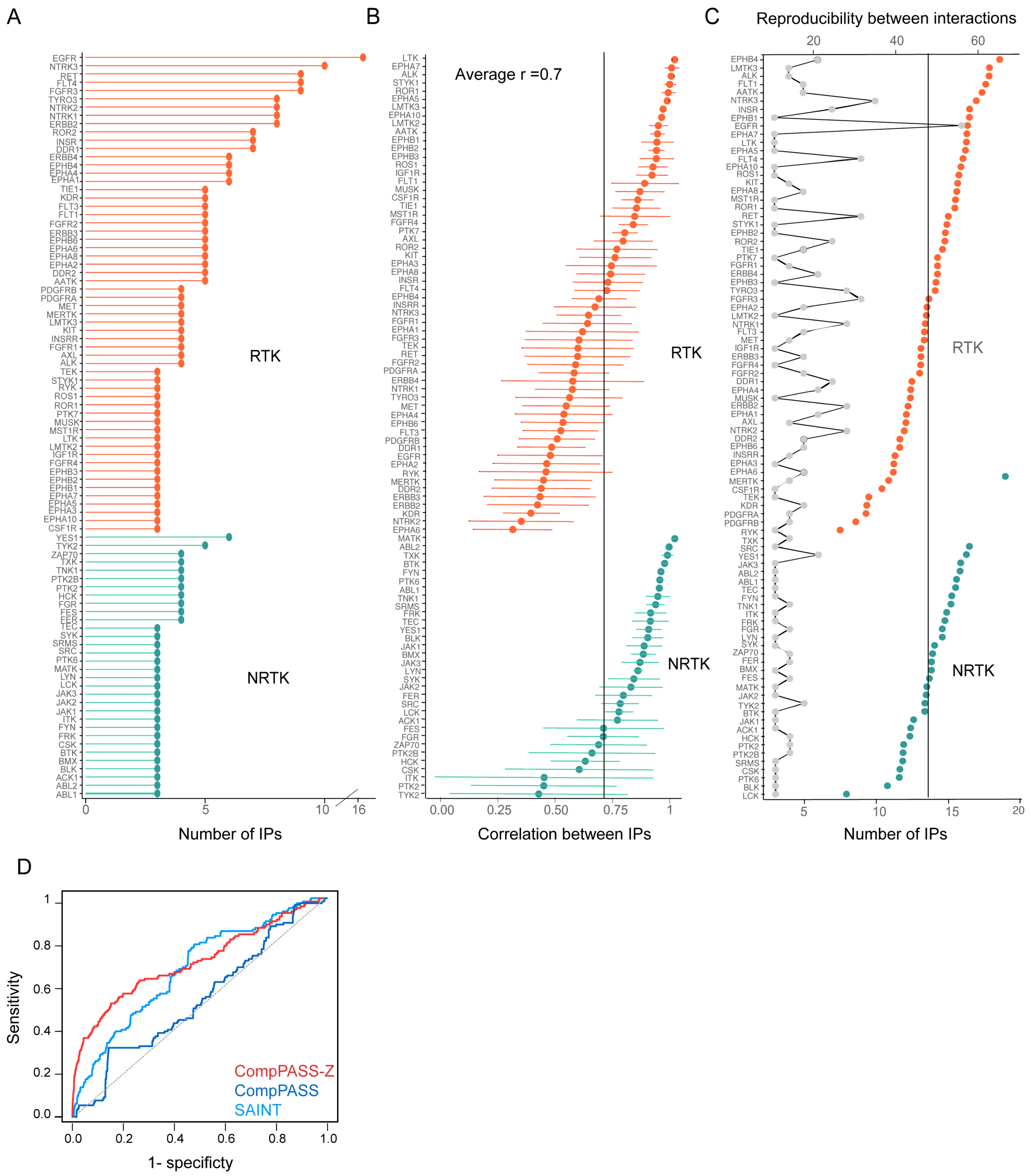


**Figure S1: Quantitative analysis of the AP-MS dataset. (A)** Number of AP-MS experiments carried out for each tyrosine kinases. **(B)** Pearson correlation of prey spectral counts among replicate experiments for each bait. Error bars indicate s.d. **(C)** Reproducibility between interactions was defined as number of preys identified in two or more replicates for the same bait (colored dots) compared with the number of AP-MS experiments carried out per tyrosine kinase bait (grey line). **(D)** ROC analysis of enrichment of MS data scored using CompPASS, CompPASS-Z and SAINT scoring methods compared with known interactions from iRefWeb database. Grey line depicts random prediction.

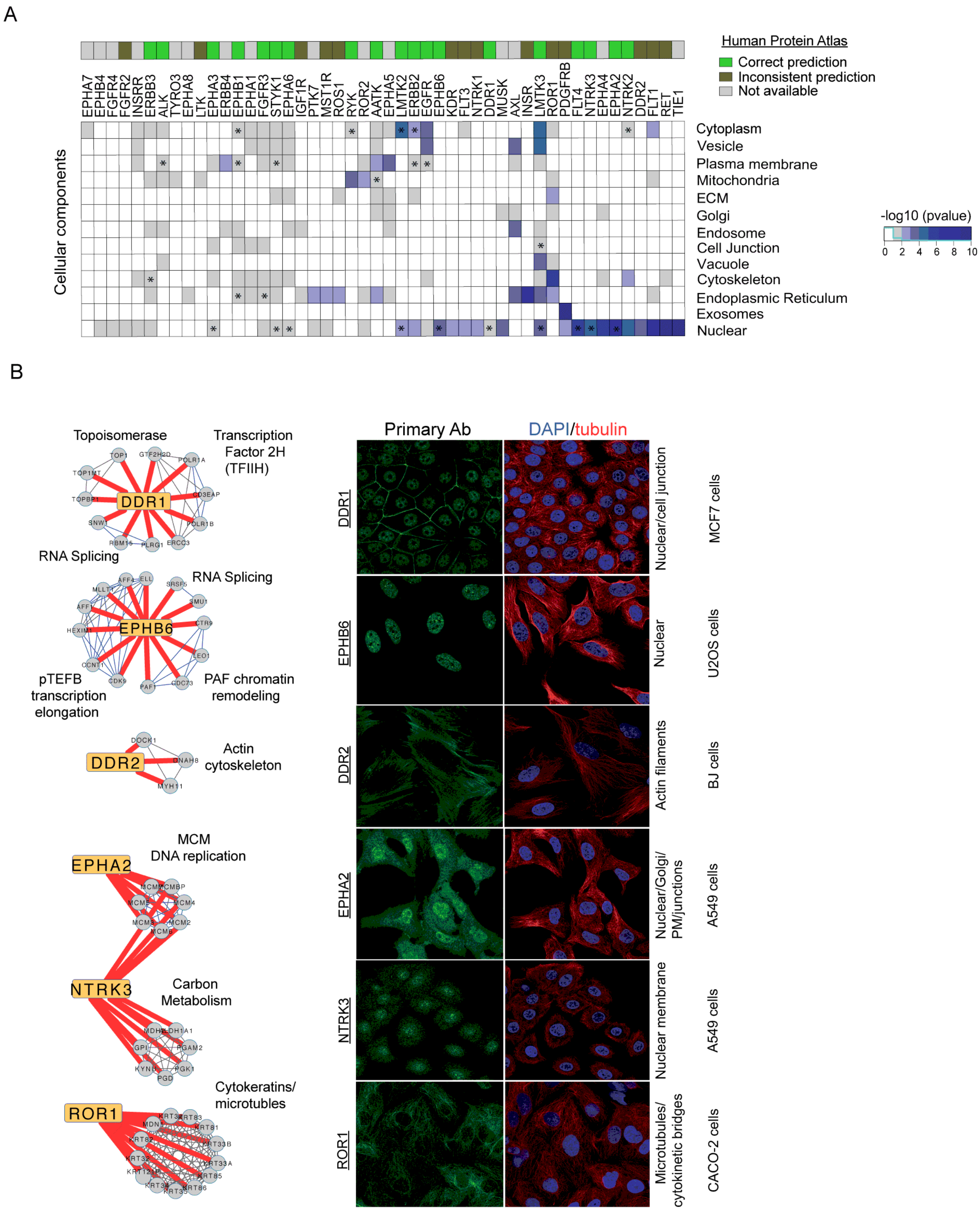


**Figure S2: Tyrosine kinase interactome predicts diverse cellular localization of receptor tyrosine kinases. (A)** Heatmap represents the predicted cellular localization of RTKs. P-values calculated using Fisher's Exact test for enrichment of GO cellular localization terms among interactors (see methods). Top row indicates the concordance with experimentally identified cellular localizations in The Human Protein Atlas database (Uhlén et al., 2005), with consistent predictions marked with an asterisk. **(B)** Network representation of interactions identified in this study (red edges) between RTKs and known pathways/complexes with known cellular localizations. Right panel shows immunofluorescence images of the indicated cancer cell line stained with primary antibodies against the listed RTK in the Human Protein Atlas. DAPI staining indicates nuclear/DNA and tubulin staining identifies microtubules.

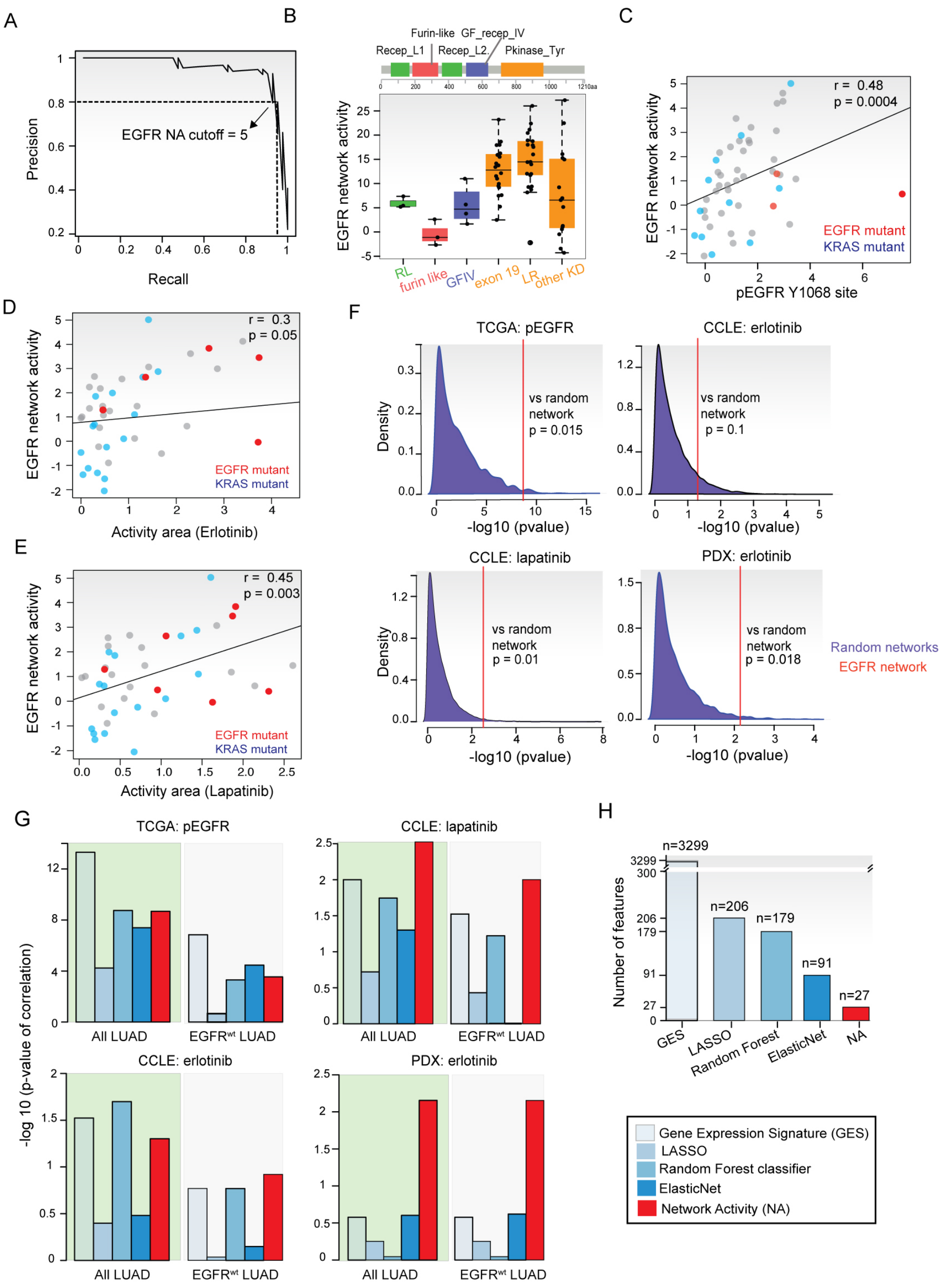


**Figure S3: EGFR network activity in lung cancer patients. (A)** Comparison of percent of samples with EGFR canonical mutations (true positives, n=42) versus KRAS mutations (false positives, n=152) at varying EGFR network activity (NA) cutoffs. A cutoff for NA score was optimized at a recall of 95% corresponding to a precision of 80%. **(B)** EGFR network activity in TCGA samples with mutations in different domains of EGFR. Median with box representing the interquartile range and whiskers 1.5x the interquartile range. **(C)** pEGFR Y1068 levels in NSCLC cell lines annotated by EGFR or KRAS mutation. EGFR network activity in NSCLC lines compared with erlotinib **(D)** and lapatinib **(E)** sensitivity measured through activity area. **(F)** Comparison of EGFR network activity with random networks composed of 27 genes whose expression was also differentially expressed in the presence of an EGFR mutation. Performance of 10,000 such random versus the real network in predicting pEGFR in the TCGA, erlotinib and lapatinib response in NSCLC cell lines and erlotinib response in NSCLC PDX models. **(G)** Comparison of EGFR NA with classical machine-learning methods. EGFR NA based prediction of pEGFR and drug sensitivities compared with that from gene expression signatures, regression models (LASSO and elastic net) and random forest classifier. Higher bars indicate more significant correlation with feature. **(H)** Comparison of number of features (genes) used in the classifier. All correlations are Spearman with corresponding two-tailed p-value shown.

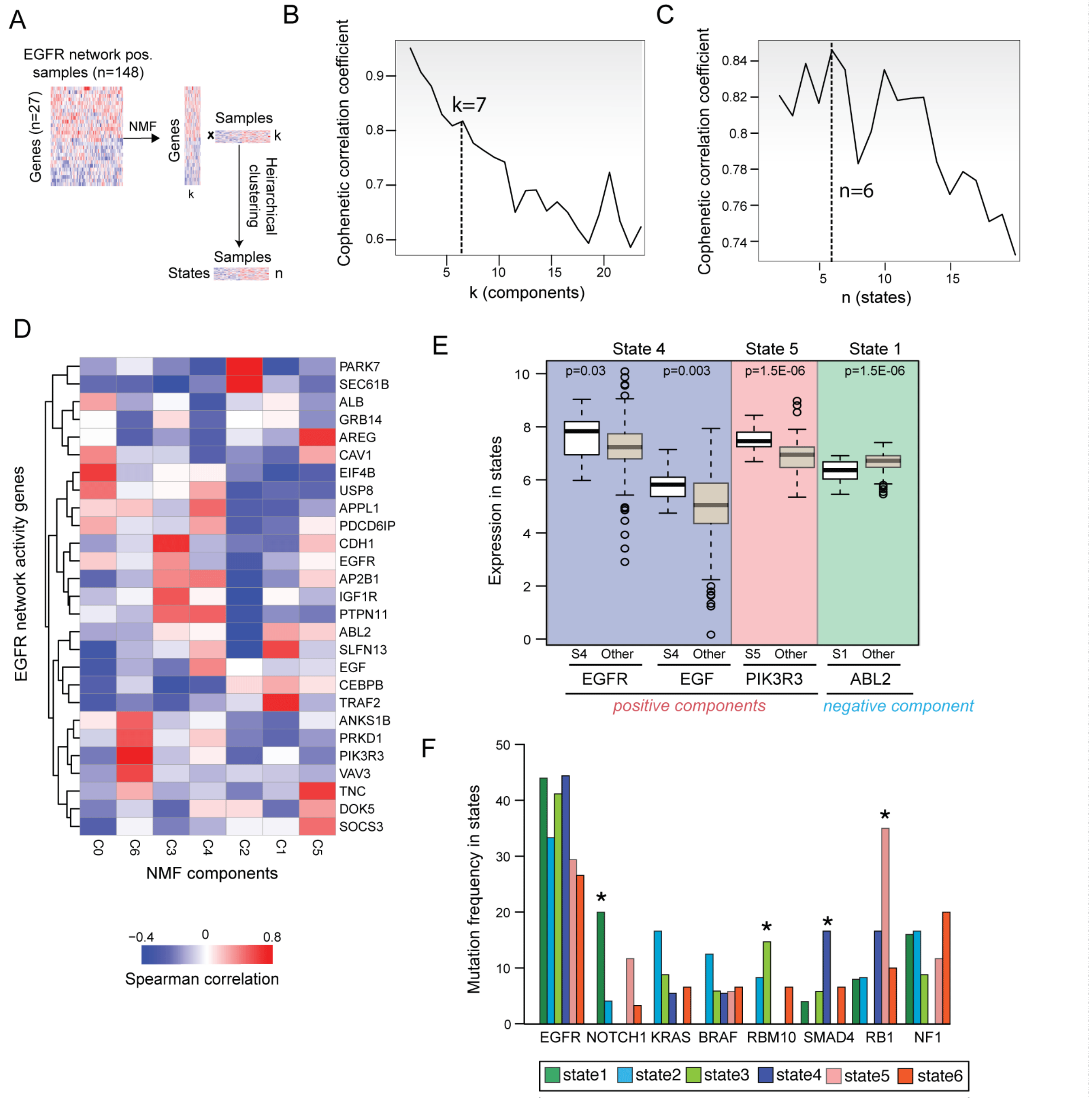


**Figure S4: NMF of EGFR network state high LUAD patients. (A)** Schematic of NMF procedure to decompose EGFR$^{n+}$ samples into components (*k*). **(B)** Identification of optimal number of components in the NMF by comparing cophenetic correlation coefficient versus number of transcriptional components (k). A peak in the cophenetic coefficient at k=7 indicates a stable decomposition with 7 distinct components. **(C)** The graph shows the cophenetic correlation coefficient as a function of different states (n). A peak in the cophenetic coefficient at n=6 indicates a stable solution with 6 states. **(D)** Spearman correlation of transcriptional components (k=7) with expression of EGFR network activity genes across EGFR$^{n+}$ samples. **(E)** Expression of selected EGFR interactors in states in which they are significantly up/down regulated compared to other EGFR$^{n+}$ samples. P-values based on Mann-Whitney test. **(F)** Mutation enrichment analysis in distinct EGFR network states of EGFR$^{n+}$ samples. For samples in a given state the frequency of mutation in known cancer genes (Lawrence et al., 2014) is shown. *=p<0.01 by Fisher's exact test.

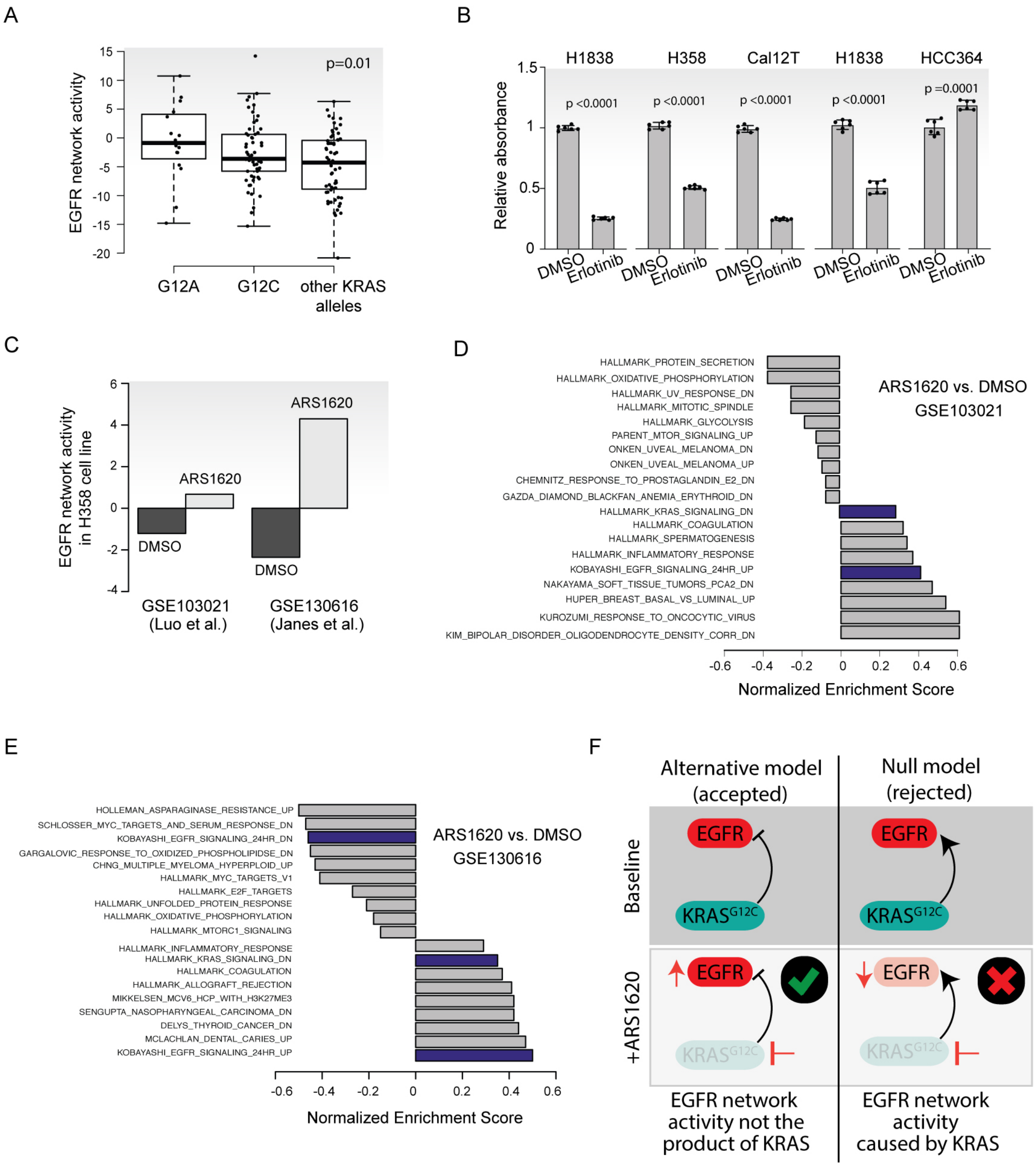


**Figure S5: Weak oncogenic mutations co-occur with EGFR$^{n+}$ patients. (A)** Comparison of EGFR network activity in the G12A and G12C mutant samples as well as samples with other KRAS mutations. P-value calculated using one-way anova. **(B)** Relative absorbance of NSCLC cell lines harboring weak oncogenic mutations upon treatment with erlotinib (1µm)**.** P-values were calculated using two-sided t-test. **(C)** Comparison of EGFR network activity before and after KRAS G12C inhibitor (ARS1620) treatment for 24 hours. EGFR network activity was calculated using RNA-Seq data deposited in GSE103021 and GSE130616 in the NCBI GEO database. **(D-E)** Gene Set Enrichment Analysis showing top perturbed gene sets following KRAS G12C inhibitor (ARS1620) treatment compared to DMSO in GSE103021 **(D)** and GSE130616 **(E)**. **(F)** Model indicating that EGFR network activation is not the product of KRAS signaling. Data indicate that KRAS suppression by ARS1620 results in the hyperactivation of the EGFR network consistent with the alternative model rather than the null.
